# Topical application of carbon dots and mesoporous silica nanoparticle-derived dsRNA-nanocomposites for the control of beet curly top virus and turnip mosaic virus

**DOI:** 10.1101/2024.12.29.628607

**Authors:** Sara Zarrabi, Carmen Rangel, Emanuel Martínez-Campos, Josemaría Delgado-Martín, Ayyoob Arpanaei, Masoud Shams-bakhsh, Leonardo Velasco

**Affiliations:** Plant Pathology Department, Faculty of Agriculture, Tarbiat Modares University, Tehran, Iran; Laboratorio de Fitopatología, Instituto Andaluz de Investigación y Formación Agraria (IFAPA), Churriana, Málaga, Spain; Scion, Private Bag 3020, Rotorua 3046, New Zealand

**Keywords:** dsRNA, carbon dots, mesoporous silica nanoparticles, SIGS, virus silencing, symptoms, turnip mosaic virus, beet curly top virus

## Abstract

The management of emerging phytoviruses in current agriculture confronts many challenges, including the appearance of new strains or the arrival of new species, requiring a multidisciplinary approach. The usual methods for the control of these pathogens are based on management practices, vector control, seed control, etc., and the introgression of genetic resistance in cultivars. But the development of resistances by classical genetic methods is a costly and time-consuming process. A promising alternative is the control of these pathogens by means of the so-called SIGS (Spray-Induced Gene Silencing) that consists of topical treatments with specific molecules derived from the viruses as double-stranded RNA (dsRNA), triggering the plant defense mechanisms. In this work, we show the control of two viruses, the potyvirus turnip mosaic virus (TuMV) and the curtovirus beet curly top virus (BCTV) by dsRNA nanocomposites with carbon dots or mesoporous silica nanoparticles in *Nicotiana benthamiana*. In the case of TuMV, the disease was significantly reduced in terms of plant photosynthetic capacity and viral titers, in the dsRNA treated plants. Moreover, when the treatments were carried out with dsRNAs as nanocomposites, the differences were even more noticeable with respect to untreated inoculated plants. In the case of BCTV, a significant delay in symptoms appearance was observed after the treatments with the dsRNA nanocomposites, but not with the naked dsRNAs. Viral titers were reduced either with the naked or the nanoparticle-delivered dsRNAs. The increased efficiency of dsRNA for virus control when supplied with the nanoparticles can be related with the enhanced dsRNA delivery reported in this work.

## Introduction

Global agriculture faces unprecedented challenges in sustaining crop productivity due to the growing pressure from pests and diseases (Savary et al. 2019). These biotic stresses are responsible for significant yield losses across staple crops, threatening food security for an increasing global population (Oerke 2006). Traditional pest and disease control methods, such as chemical pesticides and genetic resistance through breeding, are becoming increasingly ineffective due to the rapid adaptation and evolution of pathogens, as well as growing concerns about environmental safety and the persistence of chemical residues in food systems (Lamichhane et al. 2016). There is an urgent need for innovative, sustainable, and environmentally friendly alternatives that can mitigate these threats while preserving biodiversity and human health (Pretty et al. 2018; Velasco et al. 2020).

Spray-Induced Gene Silencing (SIGS) has emerged as one such promising strategy (Dalakouras et al., 2016). SIGS leverages the plant’s natural RNA interference (RNAi) mechanisms to target specific viral, fungal, or insect pathogens (Niu et al. 2021). This method involves the exogenous application of double-stranded RNA (dsRNA), designed to match sequences of the target organism’s genome (Koch et al. 2016). Upon entry into the plant, the dsRNA is processed into small interfering RNAs (siRNAs) that guide the plant’s defense machinery to degrade the corresponding viral RNA, thus silencing the pathogen’s replication and spread (Guo et al. 2016). SIGS offers the advantages of specificity, avoiding the off-target effects seen with traditional chemical pesticides, and it can be rapidly deployed without the long timelines required for developing transgenic crops or conventional breeding (Puyam and Kaur 2020; Taliansky et al. 2021). However, despite its potential, the practical application of SIGS faces several challenges. Chief among these is the instability of dsRNA in the environment (Mitter et al. 2017a). When applied to plant surfaces, naked dsRNA is highly susceptible to degradation by environmental factors, including ultraviolet (UV) light, nucleases, and temperature fluctuations (Dubelman et al. 2014). This short half-life drastically limits its effectiveness in field conditions. Furthermore, the efficiency of dsRNA uptake by plant cells is variable, and successful internalization is essential for initiating RNAi and achieving pathogen control (Dalakouras et al. 2016). These barriers have spurred interest in the use of nanotechnology to improve the delivery and stability of dsRNA.

Nanotechnology, a field that deals with the manipulation of matter on an atomic, molecular, or supramolecular scale, has made significant strides in agriculture, particularly in the development of novel delivery systems for agrochemicals and biological agents (Fraceto et al. 2016; Wang et al. 2016; Rodrigues et al. 2017). By incorporating nucleic acids into NP-delivery systems, some of the limitations associated with the direct application of nucleic acids in plants could be overcome (Choudhury et al., 2019) (Kwak et al. 2019; Mat Jalaluddin et al. 2023). NPs can protect nucleic acids from environmental degradation, enhance their cellular uptake, and enable controlled release, thereby increasing their effectiveness (Mitter et al. 2017a, b; Demirer et al. 2019). Moreover, NPs facilitate the cellular uptake of biomolecules by providing a surface charge that enhances the interaction with the plant cell membrane (Cunningham et al. 2018). This is particularly important because plant cell walls are a major barrier to the internalization of nucleic acids. Furthermore, NPs ratios can be arranged to release their cargo in a controlled manner, ensuring that the nucleic acids (e.g. DNA) are available for a prolonged period, thereby enabling the efficiency of the process (Demirer et al. 2019).

Carbon dots are a class of carbon-based nanomaterials with unique optical properties, high water solubility, and low toxicity (Das et al. 2018), making them suitable for agricultural applications such as seed priming, photosynthetic enhancers, plant stress ameliorators, sensors and delivery agents (Maholiya et al. 2023). Their small size and surface functionalization capabilities allow them to penetrate plant tissues effectively and deliver dsRNA molecules to the intracellular environment (Schwartz et al. 2020; Delgado-Martín et al. 2022a). Mesoporous silica nanoparticles, which feature tunable pore sizes and large surface areas, have been widely used for drug delivery in biomedical fields (Liang et al., 2019) (Zhou et al. 2018; Liu et al. 2021), present low cytotoxicity (Sangwan et al. 2023), and are now being adapted for agricultural purposes for pesticide (Xhu et al., 2021) or siRNA delivery in plants (Sangwan et al. 2023; Cai et al. 2024). In particular, amine functionalization of MSNs and control of pore size have allowed the controlled adsorption and release of siRNA (Steinbacher and Landry 2014).

In this study, we aimed to harness the benefits of SIGS and nanotechnology for the control of two important plant viruses: turnip mosaic virus (TuMV) and beet curly top virus (BCTV). Both viruses cause substantial economic damage to a variety of crops (Tomlinson 1987; Chen et al. 2010). TuMV, a member of the *Potyvirus* genus, has a broad host range of over 300 plant species, including economically important vegetable crops, ornamentals, and weeds. Importantly, TuMV, unlike other potyviruses, is capable of infecting brassicas (Walsh and Jenner 2002). Its genome consists of a positive-sense single-stranded RNA that encodes multiple proteins essential for its replication and virulence (Ohshima et al. 2002). The virus is transmitted by aphids in a non-persistent manner, making it difficult to control through traditional vector management strategies (Shattuck 1992). Additionally, TuMV exhibits high genetic diversity due to frequent recombination events, further complicating the development of resistant crop varieties (Walsh and Jenner 2002; Ohshima et al. 2007). BCTV, a *Be*c*urtovirus*, poses a serious threat to sugar beet production, leading to stunted growth, leaf curling, and yield losses (Strausbaugh et al. 2008). This DNA virus is transmitted by the beet leafhopper, and once inside the plant, it replicates in the phloem tissue (Soto and Gilbertson 2003). Both the severe strain of BCTV (BCTV-Svr) and beet curly top Iran virus (BCTIV) are the causal agents of beet curly top disease in Iran where they are widespread (Majidi et al. 2017). Controlling these viruses is challenging due to their efficient insect transmission, which complicates traditional management strategies such as chemical insecticides or the breeding of resistant cultivars (Bennett 1971; Lecoq and Desbiez 2012). Previous research has revealed that *N. benthamiana* and *Arabidopsis* transgenic plants for the coat protein (CP) gene of TuMV exhibited reduced or delayed symptom expression (Jan et al. 1999; Nomura et al. 2004). In another approach, workers reported that BCTV-infected *N. benthamiana* plants carrying a transgenic subgenomic viral segment showed symptom relief and a decrease in viral load (Frischmuth and Stanley 1994). Next-generation genome editing techniques offer alternatives for seeking plant virus resistance (Khan et al. 2022). *Agrobacterium* containing engineered tobacco rattle virus (TRV) that contained BCTV-specific sgRNA sequences, was infiltrated into constitutively Cas9-expressing *N. benthamiana* plants, that exhibited significantly reduced symptoms or lower viral DNA titers when infected (Ali et al. 2015). Unfortunately, these technologies currently face regulatory restrictions in Europe and other countries (NASEM, 2016).

SIGS offers a potential solution by enabling the specific silencing of viral genes involved in replication and symptom development (Kumar et al., 2014). Although the role of SIGS in plant virus control has been intensively explored, the delivery method of dsRNA remains a bottleneck (Mitter et al. 2017b). Without an efficient delivery system, the amount of dsRNA reaching the plant cells may be limited to trigger the RNAi pathway effectively (Dalakouras et al. 2016). In a previous work, we reported that the control of cucumber green mottle mosaic virus (CGMMV), an RNA virus, by SIGS, applied with an airbrush, was effective under greenhouse conditions (Delgado-Martín et al. 2022b). This treatment, however, was not effective under our conditions for a DNA virus, tomato leaf curl New Delhi virus (ToLCNDV), although recently efficacy has been reported for the treatment of naked dsRNA against ToLCNDV applied in large quantities in a similar way to mechanical virus inoculation of plants (Frascati et al. 2024).

While dsRNAs have been shown to be effective against RNA viruses, their application to DNA viruses like BCTV is less explored (Tenllado and Díaz-Ruíz 2001). In this study, we have investigated whether dsRNAs, in the form of CDs- or MSNs- nanoparticle composites, targeting the viral genes can inhibit BCTV-Svr and TuMV infection and limit disease severity by enhancing their delivery and stability in plants. Both types of nanoparticles have been shown to successfully deliver nucleic acids into plant cells (Delgado-Martín et al. 2022b; Xu et al. 2023). The CDs used in our experiments were synthesized using a hydrothermal method, and their size and surface charge were optimized to maximize dsRNA loading and release (Delgado-Martín et al. 2022a). As an initial approach, we investigated how dsRNA:NP ratios affected the entry and movement of dsRNAs within plants. Once the optimal ratios were established, these nanocomposites were applied to *Nicotiana benthamiana* plants, a model species for studying viral infection and RNAi mechanisms (Goodin et al. 2008). Our results showed that dsRNA delivered through NPs not only enhanced the stability of dsRNA but also significantly improved its uptake and silencing efficiency compared to naked dsRNA. In plants infected with TuMV, the nanoparticle-delivered dsRNA resulted in a significant reduction in viral titers, delayed the onset of symptom, and improved plant photosynthesis. Similarly, in the case of BCTV, the nanoparticle-delivered dsRNA reduced viral load and delayed the appearance of disease symptoms.

## 2. Material and Methods

### 2.1. Plants and Virus sources

Seeds of cucumber (*Cucumis sativus* cv. Bellpuig) were purchased from Semillas Fitó (Barcelona, Spain) and *N. benthamiana* seeds were already available in the lab. Virus inocula were obtained from different sources: the infectious clone p35tunos (Sánchez et al. 1998) was used as the source of the TuMV isolate UK1 (Walsh, 1989; Acc. No. AB194802) to inoculate *N. benthamiana* plants that were used as source of inoculum in the experiments. The infectious clone BCTV-Svr (including the full BCTV-Svr genome, Acc. No. X97203) originally collected from sugar beet plants in Iran was kindly provided by Dr. Behjatnia, University of Shiraz, Shiraz, Iran (Briddon et al. 1998). For the agroinoculations, *Agrobacterium tumefaciens* harboring the binary plasmids was cultured overnight in 25 ml of LB medium containing kanamycin (50 μg/ml) and rifampicin (10 μg/ml). When the OD of the culture reached 1, the bacteria were precipitated and the bacterial sediment was spread in 25 ml of sterile distilled water and acetosyringone (Sigma) was added to it at a final concentration of 200 μM. The bacteria were stirred for one hour at room temperature and 180 rpm. Two leaves with expanded leaflets from each plant were agroinoculated in the abaxial side using an insulin syringe. The TuMV isolate DSMZ was originally a field isolate from lettuce kindly provided by S. Winter and W. Menzel (DSMZ, Germany; Acc. No. MZ405633). This isolate was provided freeze-dried and was mechanically inoculated in *N. benthamiana* using the DSMZ buffer (0.05 M sodium/potassium phosphate pH 7.0, 1 mM EDTA, 5 mM DIECA, 5 mM thioglycolic acid). For the inoculation experiments described in this work, the TuMV isolates were mechanically inoculated in the *N. benthamiana* plants using the infected plants as the source. Plants were kept in the growth room at 22 °C and 16 h/8 h light/dark cycles as preliminary observations showed us that TuMV symptoms were better observed at that temperature rather than 25 °C.

### 2.2. Obtention of the constructs for dsRNA production

Constructs for the synthesis of dsRNAs were derived from the TuMV UK1 and DSMZ isolates and BCTV-Svr. The dsRNAs were produced in *Escherichia coli* HT115(DE3) cells that carried L4440-derived plasmids harboring T7 promoters at both sides of the virus gene segments. For obtaining the constructs, RNA extractions of plants infected either with the TuMV isolates UK1 or DSMZ using Trizol (Invitrogen) following the manufacture’s recommendations. The quality of the RNAs was estimated by gel electrophoresis and the quantity with the Nanodrop-2000 spectrophotometer (Thermo Fisher). Next, the cDNA synthesis was carried out using the MultiScribe Reverse Transcriptase (Applied Biosystems) in the presence of 1 mM dNTPs and RNase inhibitor for 1 h at 42°C. The cDNAs were used as templates for amplification by PCR of the viral gene segments to be cloned in vector L4440gtwy (G. Caldwell: Addgene plasmid # 11,344). Both for TuMV UK1 and DSMZ, two constructs each were designed for targeting the respective viruses isolates. For the TuMV UK1 one was derived from a segment of 510 bp of the *HC-Pro* gene within positions 2,066-2,575, rendering the dsHCPro. The other construct was derived from a 612 bp section of the *CP* gene in positions 8,854-9,465, for generating the dsCP. The amplicons were obtained from the cDNA of an infected plant using the sets of primers disclosed in Supp. Table 1). These primers included a region overlapping vector L4440gtwy for InFusion cloning (Takara). For that, plasmid L4440gtwy was digested with restriction enzymes *Hin*dIII and *Bgl*II and purified. Next, the assembly reaction was carried out using the digested vector and the respective HCPro and CP amplicons. The products of the reactions were used to transform *E. coli* Top10. Plasmids were extracted and checked by PCR, restriction analysis and sequencing and used to transform *E. coli* HT115(DE3). Similar plasmids in equivalent genomic regions were obtained for the TuMV DSMZ isolate. For the targeting of BCTV-Svr, a chimeric construct of 342 bp was obtained that included a 120 bp fragment covering parts of *Rep* (*C1*) and *TrAP* (*C2*) genes from BCTV-Svr genome fused to a 222 bp fragment covering parts of *CP* (*V1*) and *MP* (*V2*) genes from BCTIV genome (Montazeri et al. 2024). The construct was cloned into L4440 after RE digestion and the resulting plasmid was used to transform *E. coli* HT115(DE3). For the dsRNA delivery experiments in cucumber (dsNbCHE), an L4440gtwy-derived plasmid that included a 550-bp segment of the *N. benthamiana* magnesium chelatase subunit h gene (Acc. No. XM_004149349) was obtained by Gateway (Thermo Fisher) cloning, following manufacture’s protocols. The dsRNAs were obtained from the transformed *E. coli* cells grown in LB media supplemented with 6.25 g/L of lactose as inducer, 1 mM MgSO_4_, and carbenicillin (100 µg/mL) for 8 h. at 37 °C. The (ds)RNAs were extracted using Trizol as previously described and were quantified by densitometry analysis after agarose gel electrophoresis (Delgado-Martín and Velasco 2021).

### 2.3. Nanoparticles’ obtention and characterization

Synthesis of CDs was done according to Delgado-Martín et al. (2022a). Briefly, 2 g of branched polyethylenimines (bPEI, 10000 Da) dissolved thoroughly in a solution of 2 g glucose in 10 ml MilliQ water. Then, the solution was sealed in a stainless-steel autoclave lined with Teflon (100 ml capacity) and heated at 210 °C for 8 hours. When the autoclave was cooled down by remaining in room temperature, the solution was taken out and purified using a 0.22 μm filter (Millipore, Merck, Darmstadt, Germany). The obtained filtrates were dialyzed using 1000 Da molecular weight cut-off dialysis bags (Spectra/Por, Fisher, MA, USA) against 40 ml of MilliQ water shaking for 24 hours at 85 rpm and room temperature. Next, the dialysis bag was moved to a glass beaker containing 2000 ml double-deionized water and stirred overnight. Then, the water in the beaker renewed and the previous step repeated for 6 hours. Finally, the nanoparticles inside the dialysis bag were taken, lyophilized, weighted, and used for subsequent characterizations and applications. The CDs were resuspended in MilliQ water for the obtention of the dsRNA composites. MSNs were synthesized using our previously reported protocol based on the template removing approach with some modifications (Taebnia 2015). For that, 100 mg CTAB was dissolved in 48.65 ml deionized water and then stirred at 14000 rpm. As soon as the solution was clear, 350 μl NaOH (2 M) was added to the solution in a dropwise manner and then digital stirrer temperature was set on 138 °C. When the solution temperature reached 80 °C, 1 ml of TEOS (98%) was added dropwise, the lid was closed thoroughly and constant stirring was maintained for 2 hours. To remove the CTAB, the NPs were dispersed in an acidic alcoholic solution of HCl (37%) : ethanol (99%) (1:10 ratio) and incubated at 65 °C with constant stirring at 1000 rpm for 16 hours. After 16 hours, the NPs were collected by centrifugation and the previous step repeated for 3 hours. The synthesized Bare-MSNs (BMSNs) were washed three times with absolute ethanol and subsequently three times with deionized water. In the next step, the amine functionalization was carried out by adding 200 μl deionized water and 100 μl glacial acetic acid to 50 mg BMSNs dispersed in 4.6 ml absolute ethanol and stirred at 1000 rpm. Afterwards, 100 μl EDS was added to the stirring solution and the condition was kept for at least one hour. The resulting AMSNs were washed three times with absolute ethanol and deionized water. To add carboxyl groups to the AMSNs, 200 mg of succinic anhydride (SA) dissolved in 20 ml of DMF and solution was stirred for 20 minutes under a stream of nitrogen gas to evacuate the oxygen. Then, 100 mg of AMSNs that were previously well-dispersed in dimethylformamide (DMF) were added to the SA solution in a dropwise manner. The lid was sealed completely and the mixture was stirred overnight. The carboxyl-functionalized MSNs (CMSNs) were washed three times with absolute ethanol and deionized water. In the last step, positively charged MSNs prepared by coating CMSNs with polyethyleneimine (PEI) (PMSNs). First, CMSNs were precipitated in microtubes and redispersed in 1.5 ml of phosphate-buffer saline (PBS, 10 mM, pH 7.4) containing NHS (0.2 mg/ml) and EDC (1 mg/ml) and were manually shaken for about 5 min. The nanoparticles were immediately centrifuged for 3 min at 13000 rpm. Then, each pellet was dispersed in 1.5 ml of PBS containing PEI (2 mg/ml) and the microtubes were centrifuged at 15 rpm for 1 hour at room temperature. Finally, the obtained PMSNs were washed three times with PBS (10 mM) and deionized water. Transmission electron microscopy (TEM) (Thermofisher, FEI Talos F200X) was used for studying the size and morphology of the MSNs dispersed in deionized water (40 μg/ml) and a population was used to obtain the average size with the ImageJ software. The hydrodynamic diameter and Zeta (ζ) potential of MSNs (40 μg/ml) were measured using the Zetasizer (Zetasizer Nano ZS, Malvern, UK). Surface functionalization of each step (B-, A-, C-, and P-MSNs) was evaluated on 5 mg of lyophilized MSNs using Fourier-transform infrared (FT-IR) analysis of the spectra collected with the Tensor 27 spectrophotometer (Bruker, Bremen, Germany) using a Gate Single Reflection Diamond ATR System accessory. A standard spectral resolution of 4 cm^−1^ in the spectral range 4000–400 cm^−1^ and 64 accumulations were used.

### 2.4. Characterization of the dsRNA:PMSNs composites

PMSNs were well dispersed using a sonicator bath with medium intensity for 3 minutes prior to adding the dsRNAs. To determine the optimum dsRNA:PMSNs ratio, 2.5 μg of dsRNAs derived from the *N. benthamiana* magnesium chelatase (dsNbCHE) were electrostatically loaded on PMSNs with different ratios (1:0, 0:1, 0:10, 1:1, 1:5, 1:10, 1:20, 1:30, and 1:40) in 20 μl final volume for 1 hour. The assay was replicated trice. One replicate was electrophoresed on the 2% agarose gel for visualization. For the other two replicates, the samples were centrifuged for 10 min at 6000 rpm and the amount of RNA in supernatant was measured with the Nanodrop. Loading capacity (LC) and loading efficiency percent (LE%) of each loading ratio were calculated using equations 1 and 2, respectively.

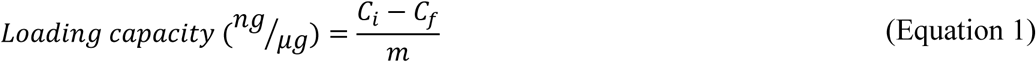

Where, *C_i_* is the initial concentration of dsRNA (ng), *C_f_* is the final concentration of dsRNA in supernatant (ng) and *m* is the PMSNs mass (μg).

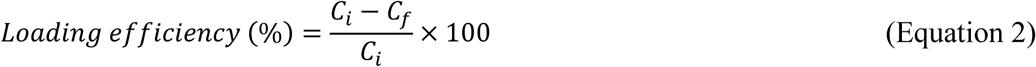

Equilibrium concentration (*C_e_*) of the loading reactions were calculated using equation 3 to plot the dsRNA adsorption isotherm on PMSNs.

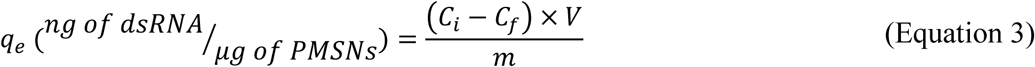

Where *V* is loading reaction volume (μl) .

Correlation of the equilibrium data with Langmuir (Equation 4), Freundlich (Equation 5), and Langmuir-Freundlich (Equation 6) models was studied by fitting the empirical adsorption isotherm on mentioned models. These isotherms relate the quantity of adsorbed material per adsorbent mass (*q_e_*) with the equilibrium concentration *C_e_* in the fluid phase.

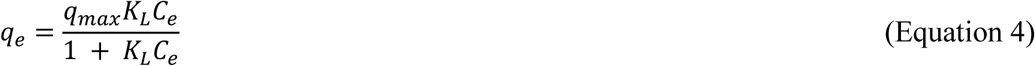

Where, *q_max_* is the maximum dsRNA adsorption capacity of PMSNs (ng/μg), *K_L_* is the Langmuir’s constant (μl/ng) and *C_e_* is the equilibrium concentration of dsRNA (ng).

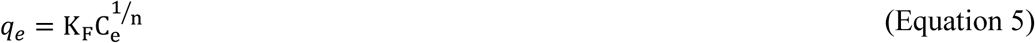

*K_F_* is the Freundlich’s constant (ng/mg), *C_e_* is the equilibrium concentration of dsRNA (ng) and *n* is the heterogeneity index.

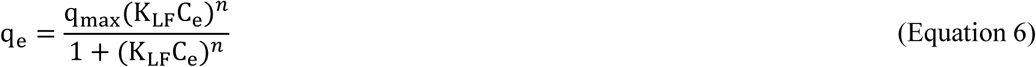

*q_max_* in the L-F isotherm model is the maximum dsRNA adsorption capacity of PMSNs (ng/μg), *K_LF_* is the affinity constant for adsorption (L/mg), *C_e_* is the equilibrium concentration of dsRNA (ng) and *n* is the heterogeneity index. The values of the constants in the different models were estimated using the solver add-in in Microsoft Excel.

### 2.5. Release of dsRNA from PMSNs

Sodium dodecyl sulfate, SDS (0.1%, 0.2%, and 0.3%) and ethylenediaminetetraacetic acid, EDTA (5 and 10 mM) treatments on dsRNA:PMSNs used to study the dsNbCHE release from the PMSNs. For that, we used in these assays a 1:10 loading ratio in 20 μl final volume that were incubated for 30-min incubation at room temperature. Then, solutions of SDS or EDTA were added to the nanocomposite solutions and mixed by pipetting. The microtubes were next incubated at room temperature for another 30 min and the samples inspected on 2% agarose gels.

### 2.6. DsRNA delivery to plant leaves

DsRNA delivery was studied on cucumber and *N. benthamiana* leaves at 3 and 5-6 leaf stages, respectively. In all experiments of this section, each leaf was sprayed with 60 μg of (ds)RNA extract that included 3.5 μg (∼5 %) of the corresponding specific dsRNA, according to our estimations from the densitometry analyses (not shown) by using an airbrush at 2.5 bar pressure. There were no viral inoculations involved in these experiments. For sampling, the leaves were washed thoroughly with double-distilled water using a wash bottle and then a clean airbrush to wash away any surface remaining dsRNA three days post-treatment (dpt) and sampled on the middle of the leaves. In a set of experiments, to examine the different loading ratios of dsRNA:PMSNs in dsRNA delivery to plant cells, cucumber plants were used due to its larger leaf surface and ease of spraying. On each plant, at the two to three leaf stage, one leaf was sprayed with dsNbCHE and treatments were prepared as follows: dsRNA:CDs (1:0.5), dsRNA:PMSNs (1:1, 1:10, and 1:20), naked dsRNA, and as mock the dsRNA derived from the CP of TuMV DSMZ (dsTuMV-CP) with six replications per condition. The final spray volume for CDs and PMSNs were 400 μl and 2000 μl, respectively. Noteworthy, PMSNs are much bigger in size compared to CDs; hence, to avoid the precipitation of dsRNA:PMSNs nanocomposites (especially in higher loading ratios as 1:30 and 1:40) it was necessary further dilution with MilliQ water. In another round of experiments, and using three replications, dsRNA derived from BCTV (dsBCTV) was sprayed on *N. benthamiana* leaves to evaluate dsRNA presence in sprayed and unsprayed leaves. The treatments were: dsBCTV:CDs (1:0.5), ds dsBCTV:PMSNs (1:10 and 1:30), naked dsBCTV, and mock. The dsRNA:PMSNs loading ratio of 1:10 was chosen from the previous experiment and 1:30 was also tested to see what happens in the presence of excessive amount of PMSNs. For the evaluation of plant dsRNA movement after spraying with the nanocomposites we choose cucumber. For that, one side of cucumber leaves was covered with aluminum foil to avoid the spraying followed by dsRNA treatments on the other half-side. After 3 dpt, leaf samples were taken from each leaf-side and the dsRNAs were quantified as described below. In a fourth round of experiments to evaluate the stability of dsRNA in plants with time, in three replications, two leaves of each cucumber plant were sprayed with dsBCTV. Tissue samples were taken from sprayed leaves at 7 dpt. In this case, the treatments were as follows: dsBCTV:CDs (1:0.5), dsBCTV:PMSNs (1:10), naked dsBCTV, and mock as negative control.

For the quantitations, RNA extractions from leaf tissues (100 mg) were performed using Trizol followed by cDNA synthesis as described above. The subsequent qRT-PCR reactions were carried out, in two technical replicates, using diluted cDNA (1:1), Kapa SYBR Fast qPCR Master Mix (2X) Universal (Kapa Biosystems), and the corresponding forward and reverse primers for the dsNbCHE, the *C. sativus* RNA 18S gene (as endogenous reference in cucumber), the dsBCTV and the *N. benthamiana* elongation factor 1α (*NbEF-1α*), each with the final concentration adjusted to 500 nM (Supp. Table 1). The qPCR program was as follows: initial denaturing of 3 min at 95 °C, 39 cycles of 10 seconds at 95 °C denaturation, 10 seconds at 55 °C annealing and 30 seconds at 72 °C extension alongside data collection at the end of each cycle, followed by a denaturing step of 10 seconds at 95 °C and the melting program consisting of 5-second ramps from 65 to 95 °C seconds with 0.5 °C/sec increments. The geometric mean of their expression ratios was used as the normalization factor in all samples for measuring the quantification cycle (Cq). The relative expressions (fold) of the (ds)RNA amounts were compared based on the calculations done with the 2^−ΔΔCq^ method. One-way ANOVA of the relative expression differences was performed followed by mean separation using Tukey’s HSD post hoc analysis when the data distributed normally or Kruskal-Wallis when non-parametric followed by Dwass-Steel-Critchlow-Fligner two-to-one comparisons, with the software Jamovi v. 2.3.21 (https://www.jamovi.org).

### 2.7. Evaluation of TuMV and BCTV-inoculated plants after the dsRNA treatments

In the inoculation experiments in *N. benthamiana* and the corresponding treatments, six plants per condition (positive control, mock, naked dsRNA-treated, CD:dsRNA-treated or MNSP:dsRNA-treated plants) were evaluated in each of the two consecutive assays (repetitions) performed (in total 12 plants per condition). The mock used in the negative controls was unspecific dsRNA derived from the *MP* gene from CGMMV (Velasco and Delgado, 2021). When the plants were four weeks old (six leaves), they were sprayed with either naked dsRNA or nanocomposite-dsRNA using an artist airbrush at 2.5 bar following with the inoculation with the respective viruses at 3 dpt. Symptoms of TuMV-inoculated plants were scored based on the chlorophyll content as quantified with the SPAD-502Plus (Konica-Minolta). For that, on each plant three measures per leaf were taken at the upper, medium and lower leaves, and the average of 12 biological replicates (plants) per condition were considered in the statistical analyses. BCTV-induced symptoms were scored from 0 to 4 after simplifying the rating scale introduced by Montazeri et al. (2024) (Table S2). Average symptoms scores were obtained at 9, 14, 16, 18, and 23 dpi for each treatment and used to calculate the area under disease progress curve (AUDPC). Then, one-way ANOVA of the AUDPCs was performed followed by mean separation using Tukey’s HSD post hoc analysis to investigate the relationship between the disease progress and the respective treatments.

### 2.6. Virus quantitations

Total RNAs were extracted from the TuMV-inoculated plants, including the untreated and dsRNA-treated plants. For that, 100 mg of plant tissue from the leaves were extracted using Trizol (Invitrogen) following manufacturer’s recommendations. RNA quality was evaluated in agarose gels and the quantity was performed using the Nanodrop. About one microgram of total RNA was used for the reverse transcription and cDNA obtention with the High-Capacity cDNA Reverse Transcription Kit. Each qPCR reaction (20 µL final volume), in triplicate, contained 1 µL of the cDNA, 10 µL of KAPA SYBR Green qPCR mix, and 500 nM each of the forward and reverse primer (Table S1). In separate reactions, we included the primers for the *NbEF1α* gene as endogenous reference. Specificity of the amplicons obtained was checked using the Bio-Rad Optical System software v.2.1 by means of melting-curve analyses (60 s at 95 °C and 60 s at 55 °C), followed by fluorescence measurements (from 55–95 °C, with increments by 0.5 °C). The relative expressions of the RNA amounts were compared as described above. Samplings for BCTV quantitation were taken at 9- and 16-days post-inoculation (dpi) from the first and second leaves above the last inoculated leaf, respectively, and subsequently the total DNA was extracted according to Accotto et al. (2000). All samples were diluted to the final concentration of 200 ng/μl and kept for studying relative accumulation of BCTV using quantitative PCR (qPCR). The qPCR reactions contained DNA (200 ng), Kapa Master Mix and, in independent reactions, the BCTV-Svr or *NbEF-1α* (as reference gene) forward and reverse specific primers (final concentration 500 nM) and carried out in technical replicates and with a final volume of 20 μl. The qPCR was set up as follows: initial denaturing step of 3 min at 95 °C, 39 cycles of 10 seconds at 95 °C denaturation, 10 seconds at 60 °C annealing and 30 seconds at 72 °C extension alongside data collection at the end of each cycle, and the melting program setting as described above. The relative expression of viral DNAs using the Cq values was estimated as above.

## 3. Results

### 3.1 Characteristics of the nanoparticles and the dsRNA nanocomposites

Two types of nanoparticles were obtained in this study: (1) carbon dot-based nanoparticles (CDs), prepared and characterized as previously reported with some modifications (Delgado-Martín et al. 2022a), and (2) mesoporous silica nanoparticles (PMSNs), synthesized as described in the above section. An average size of 5 nm and a ζ potential of +11.94 mV were observed in the CDs (Fig. S1). TEM observation of a population of about 400 PMSN determined an average diameter of 89.07 ± 14.71 nm (Figure 1A). The size distribution histogram indicated that 86% of the nanoparticles had diameters ranging between 70-115 nm, fitting well to a normal distribution pattern (Figure 1B). Dynamic light scattering (DLS) measurements were performed to determine the hydrodynamic diameter and surface charge of the nanoparticles at each stage of functionalization (Table 1). The DLS analysis revealed remarkable changes in nanoparticle charge, confirming successful functionalization. Bare mesoporous silica nanoparticles (BMSNs) exhibited a negative charge (−20.3 mV), which shifted to a positive value (+2.95 mV) following amino-acidification (AMSNs). Further functionalization with carboxyl groups resulted in a return to a negative charge (−31.5 mV) for the carboxyl-modified silica nanoparticles (CMSNs). The final step, involving the loading of PEI polymeric groups onto CMSNs, resulted in a charge increase to +22.55 mV, indicating successful formation of PMSNs, necessitated for the loading of the nucleic acids. The resulting PMSNs also offered a good polydispersity index (PdI) (Table 1). A steady increase in hydrodynamic diameter was observed with each functionalization step. The hydrodynamic diameter of PMSNs measured by DLS was 309.1 nm, which was 3.5-fold larger than the diameter obtained from TEM images (89.07 nm) (Table 1).

**Figure 1.**
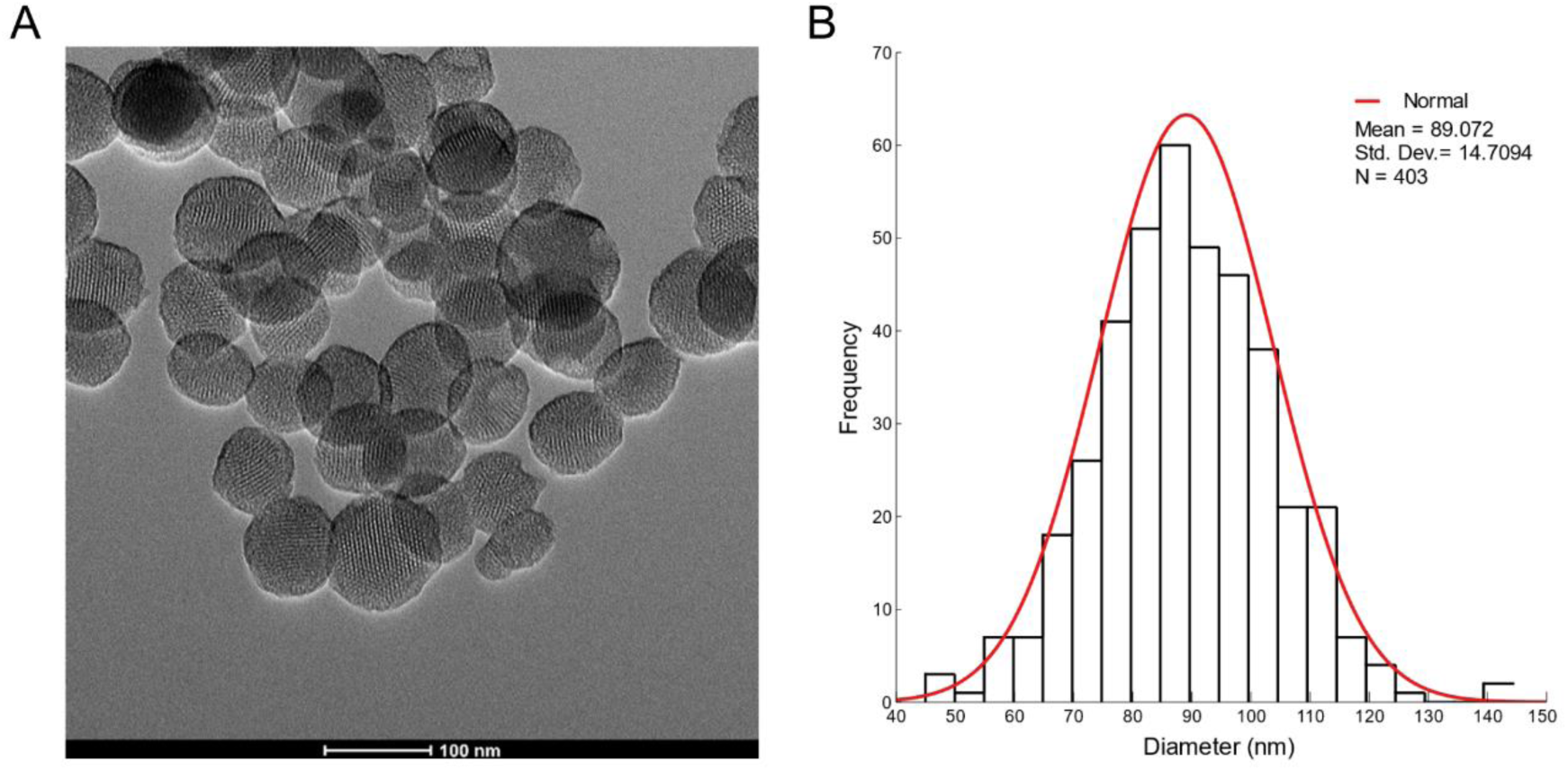
Transmission electron microscope (TEM) image of PMSNs (A) and normal distribution histogram of PMSNs nanoparticle diameter size (B).

**Table 1.** Hydrodynamic size, polydispersity index (PdI) and surface charge of the MSNs.

|  | **Zeta ($\zeta$) potential (mV)** | **Size (d.nm)** | **Pdl** |
| --- | --- | --- | --- |
| Bare MSNs (BMSNs) | -20.3 | 275.6 | 0.097 |
| Aminated MSNs (AMSNs) | +2.95 | 285.1 | 0.043 |
| Carboxylated MSNs (CMSNs) | -31.5 | 297.2 | 0.228 |
| Polyaminated MSNs (PMSNs) | +22.55 | 309.1 | 0.189 |
| dsRNA-coated PMSNs | -11.83 | 364.3 | 0.204 |

The FT-IR spectra of the MSNs confirmed the successful incorporation of functional groups at each stage of the synthesis process (Figure 2). BMSNs displayed peaks corresponding to Si-OH (821–991 cm⁻¹), Si-O-Si (1113 cm⁻¹), H₂O (1650 cm⁻¹), and O-H (3478 cm⁻¹). Upon functionalization with amino groups, the FT-IR spectrum of AMSNs exhibited peaks corresponding to N-H bonding (1580–1650 cm⁻¹), C-N-N (1659 cm⁻¹), C-H (2835–2953 cm⁻¹), and N-H stretch (3300–3500 cm⁻¹). After further modification with carboxyl groups, peaks related to COOH absorption (1413 cm⁻¹), C=O vibrations (1750–1700 cm⁻¹), and amide C=O strain (1690–1630 cm⁻¹) were observed in CMSNs. Finally, for PMSNs, the FT-IR spectrum exhibited prominent -N-H and -C-H stretching bands at 3300–3500 cm⁻¹ and 2835–2953 cm⁻¹, respectively, confirming the successful loading of PEI.

**Figure 2.**
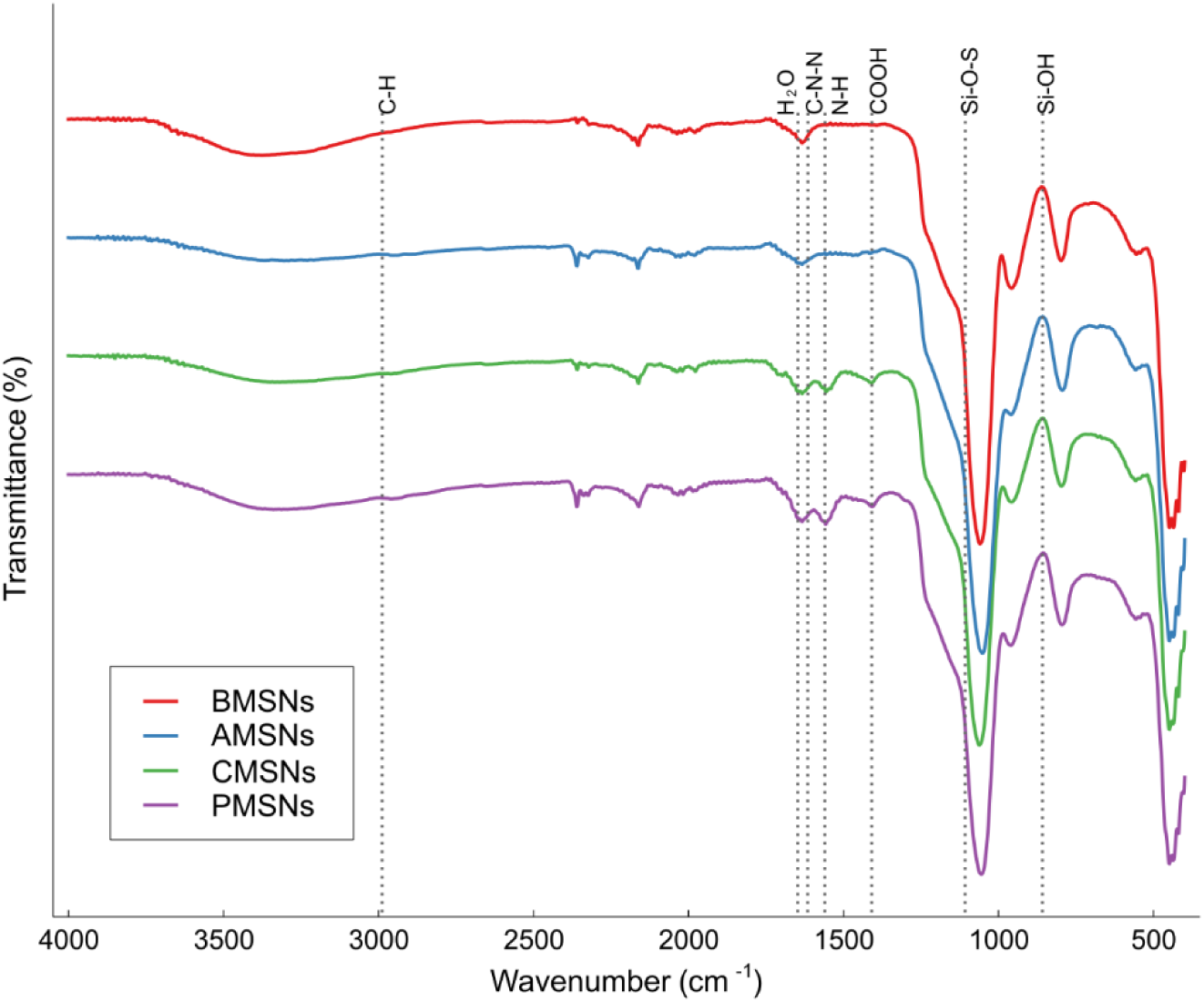
FT-IR spectra of bare silica nanoparticles (BMSNs), amine-(AMSNs), carboxylated-(CMSNs), polyamine-(PMSNs) and dsRNA:PMSNs.

### 3.2. Nanoparticle-dsRNA interactions and adsorption studies

After characterization of the nanoparticles, the interaction between CDs and PMSNs and dsRNA was investigated. Migration of dsRNA in electrophoresis was delayed when increasing the CD ratio in the dsRNA:CD composites (Figure S1). The hydrodynamic diameter of the dsRNA:PMSN (1:10 ratio) composites showed an increase with respect to pristine PMSNs (Table 1), while keeping a closer value of the PdI. In addition, dsRNA loading capacity and efficiency were assessed by incubating at room temperature different ratios of PMSNs with a constant dsRNA amount (2.5 μg). The nanocomposites were run on agarose gels to observe the dsRNA loading behavior on PMSNs (Figure 3A). While at dsRNA:PMSNs ratios 1:1 and 1:5 the dsRNA was not fully loaded, from ratios 1:10 and above the dsRNA was not visible in the gels, indicating the absence of free dsRNA molecules and that the dsRNA was stuck in the wells in the form of complex with the PMSNs. These assays allowed the calculation of the loading capacity (LC) and loading efficiency percent (LE%) of dsRNAs on PMSNs after the quantitation of remnant RNAs in supernatants after precipitation of dsRNA:PMSNs nanocomposites (Figure 3B). Given that the amounts of PMSNs were 30 and 40 times the amount of dsRNA, the LE% reached 65.7% and 68.9%, respectively, representing more electrostatically absorbed dsRNA due to the presence of the more positive-charged PMSNs. On the other hand, at the lower loading ratios (i.e. 1:1 and 1:5), the surface of the PMSNs was saturated with the dsRNA molecules, resulting in higher LC and lower LE%. The loading capacity difference between the 1:20 and 1:10 loading ratios was significant and it did not change dramatically at loading ratios of 1:1 and 1:5. Thus, the 1:10 ratio was shown as suitable for loading dsRNA molecules on PMSNs. The isotherm derived from the binding assays was modeled using the Langmuir, Freundlich, and Langmuir-Freundlich models, with the latter model providing the best fit (*R*^2^ = 0.95) for the interaction between dsRNA and PMSNs (Figure 3C) (Table S3). Regarding the FT-IR analysis of dsRNA:PMSN, revealed that all the peaks that were observed in the absorption spectrum of the pristine PMSN sample were present. Besides, in the sample loaded with dsRNA we could detect an asymmetric PO₂⁻ stretching at 1220-1240 cm⁻¹ (Figure S2).

**Figure 3.**
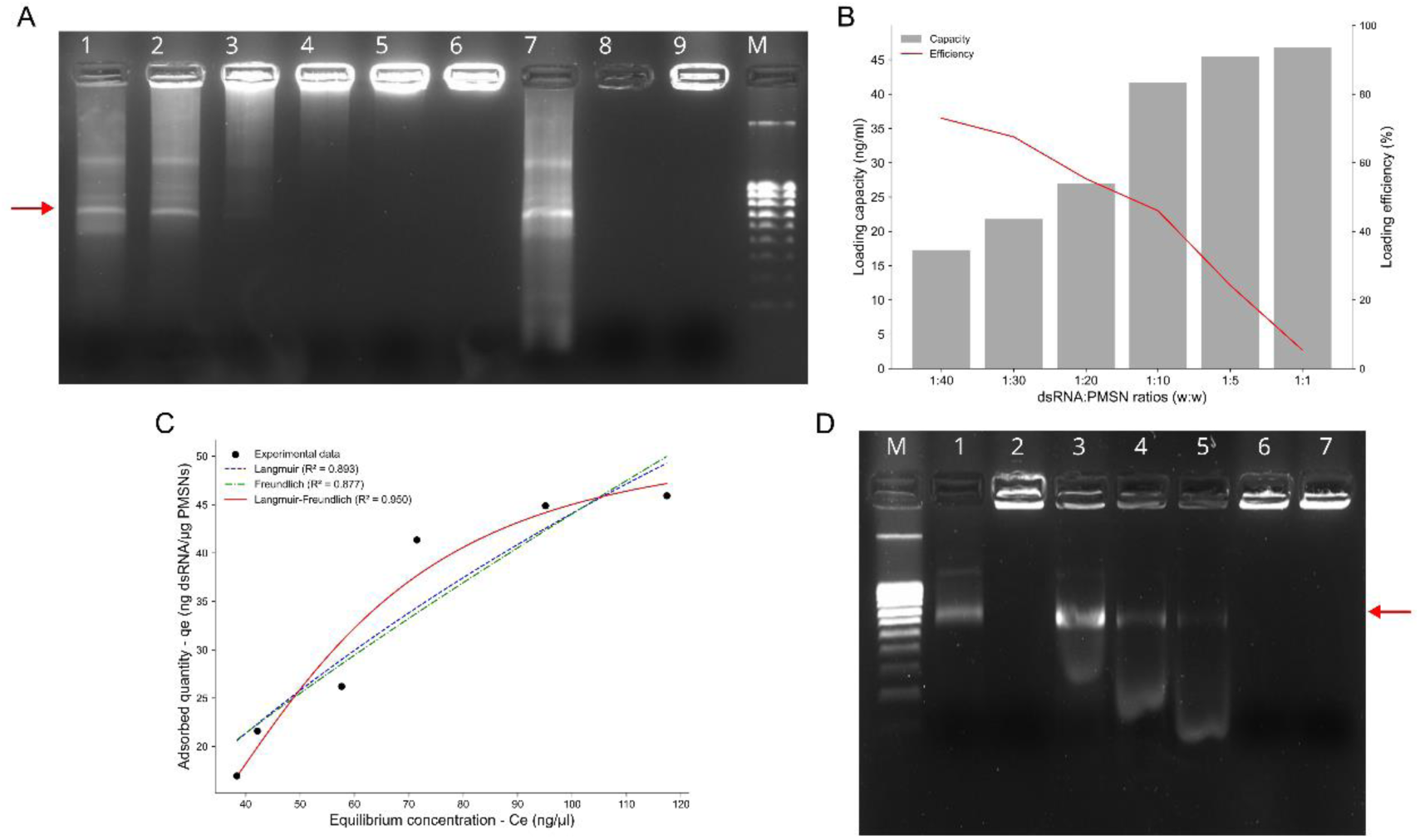
A) Retardation gel of dsRNA with increasing amounts of PMSNs on 2% agarose gel. Different wells contained different ratios of dsRNA:PMSNs pool: 1:1 (1), 1:5 (2), 1:10 (3), 1:20 (4), 1:30 (5), 1:40 (6), 1:0 (7) (negative control PMSNs), 0:1 (8), 0:10 (9), (8 and 9 dsRNA negative controls) and M: molecular weight marker (Ladder V NZY Tech). dsRNA is indicated by a red arrow. B) Loading capacity loading efficiency of dsRNA on PMSNs (loading a constant amount of 2.5 μg of dsRNA on variable amounts of PMSNs). C) Isotherm plot of dsRNA adsorption on PMSNs and fitting to the Langmuir, Freundlich and Langmuir-Freundlich models for the experimental data of dsRNA adsorption on PMSNs, being *q_e_* the quantity of adsorbed material per adsorbent mass. D) Release of dsRNA release from PMSNs on 2% agarose gel. dsRNA control (untreated) (1), PMSNs control (0:1) (2), dsRNA:PMSNs + 0.1% SDS (3), dsRNA: PMSNs + 0.2% SDS (4), dsRNA:PMSNs + 0.3% SDS (5), dsRNA:PMSNs + 5 mM EDTA (6), dsRNA:PMSNs + 10 mM EDTA (7), M: molecular weight marker (Ladder V). The dsRNA band is indicated by a red arrow.

The hydrodynamic diameter as measured with DLS of the dsRNA:PMSNs was slightly higher than pristine PMSNs suggesting the presence of dsRNA molecules on the surface of PMSNs. Besides, we observed a remarkable decrease of surface charge according to ζ potential measurements, from +22.55 (pristine PMSNs) to −9.91 mV (dsRNA:PMSNs) for 1:10 ratio confirming that the positive surface of PMSNs were coated with the negative RNA molecules via electrostatic interactions (Table 1). The dsRNA:CD composite (1:0.5 ratio) had a surface charge of −7.7 mV, whereas naked dsRNA resulted in −13.5 mV. SDS and EDTA treatments were used to study the dsRNA release from the PMSNs. SDS (0.1%), but not EDTA, successfully interfered with the electrostatic interactions among negative phosphate groups of dsRNA molecules and positive polyamine groups of PMSNs due to their amphipathic nature (Figure 3D).

### 3.3. Nanoparticle carriers improve dsRNA delivery to the plant and enhance its movement and stability

The delivery of dsRNA either naked or in the form of nanocomposites to plant leaves was assessed by qRT-PCR at 3 days post-application. For that, a regular amount of 60 μg (ds)RNA bacterial extract was applied to leaves at varying dsRNA:NP ratios. In cucumber, sprayed with dsNbCHE, the dsRNA:NP composites showed significant increases in dsRNA delivery compared to naked dsRNA (Figure 4A). Average Cq values resulted 17.8, 20.2, 18.87, 20.28 and 22.45 for the dsRNA:CD (1:0.5), dsRNA:PMSN (1:1), dsRNA:PMSN (1:10), dsRNA:PMSN (1:20) and naked dsRNA, respectively. These values resulted in 5.1 and 4.3-fold delivery increases of the dsRNA:CD and dsRNA:PMSN (1:10) with respect to naked dsRNA, respectively. In *N. benthamiana*, a similar trend was observed when spraying with dsBCTV and their nanocomposites. In this case, the average Cq values resulted 18.3, 18.9, 21.17 and 21.3 for dsRNA:CD (1:0.5), dsRNA:PMSN (1:10), dsRNA:PMSN (1:20) and naked dsRNA, respectively. The respective fold increases showed significant differences (*P* < 0.05) with respect to the naked dsRNAs were 2.43 and 3.52 for the dsRNA:CD and dsRNA:PMSN (1:10) (Figure 4B). Thus, increases in the dsRNA:PMSNs loading ratios up over 1:10 did not result in improved dsRNA uptake, rather decreased it, indicating that this ratio was optimal and chosen for the subsequent experiments. The movement of the dsNbCHE in cucumber were evaluated after spraying one half of the leaf and sampling both in the sprayed half-leaf and on the non-sprayed half, that was protected by aluminum foils. The results confirmed the results from the above experiments, but including the additional dsNbCHE:CD 1:2.5 ratio in the assays (Figure 4C). When compared with naked dsRNAs, delivery assisted the CDs in ratio 1:2.5 was the highest (16-fold) followed by the 1:0.5 ratio (5.1-fold) and the dsNbCHE:MNSP 1:10 and 1:20, 4.7 and 4.4-fold, respectively. When examining the samplings in the distal part of the leaf, the dsRNA delivered the CDs in ratio 1:05 resulted 50-fold higher than when the dsRNA was delivered naked, and 2.3-fold when delivered as dsNbCHE:MNSP 1:10 ratio (Figure 5D). For the proximal and distal half-leaves, the average Cq values when treated with dsNbCHE:CD 1:0.5 were 17.8 and 22.9. For the dsNbCHE:MNSP 1:10 were 18.9 and 28.65, in the proximal and distal half-leaves, respectively. Treatments with the naked dsRNAs showed the average Cq values 22.42 and 28.28, respectively, in the proximal and distal half-leaves. We further explored the stability with time of dsBCTV in *N. benthamiana* plant leaves after nanoparticle-mediated delivery at 7-days post-application. After this period, the Cq values resulted 20.8, 21.2 and 21.9 for dsRNA:CD (1:0.5), dsRNA:PMSN (1:10) and naked dsRNA, respectively. From these values, it resulted that the dsRNA:CD treatments showed a 2.1-fold increase, while dsRNA:PMSN (1:10) treatments exhibited a 1.8-fold increase compared to the naked dsRNA treatments (Figure 4E).

**Figure 4.**
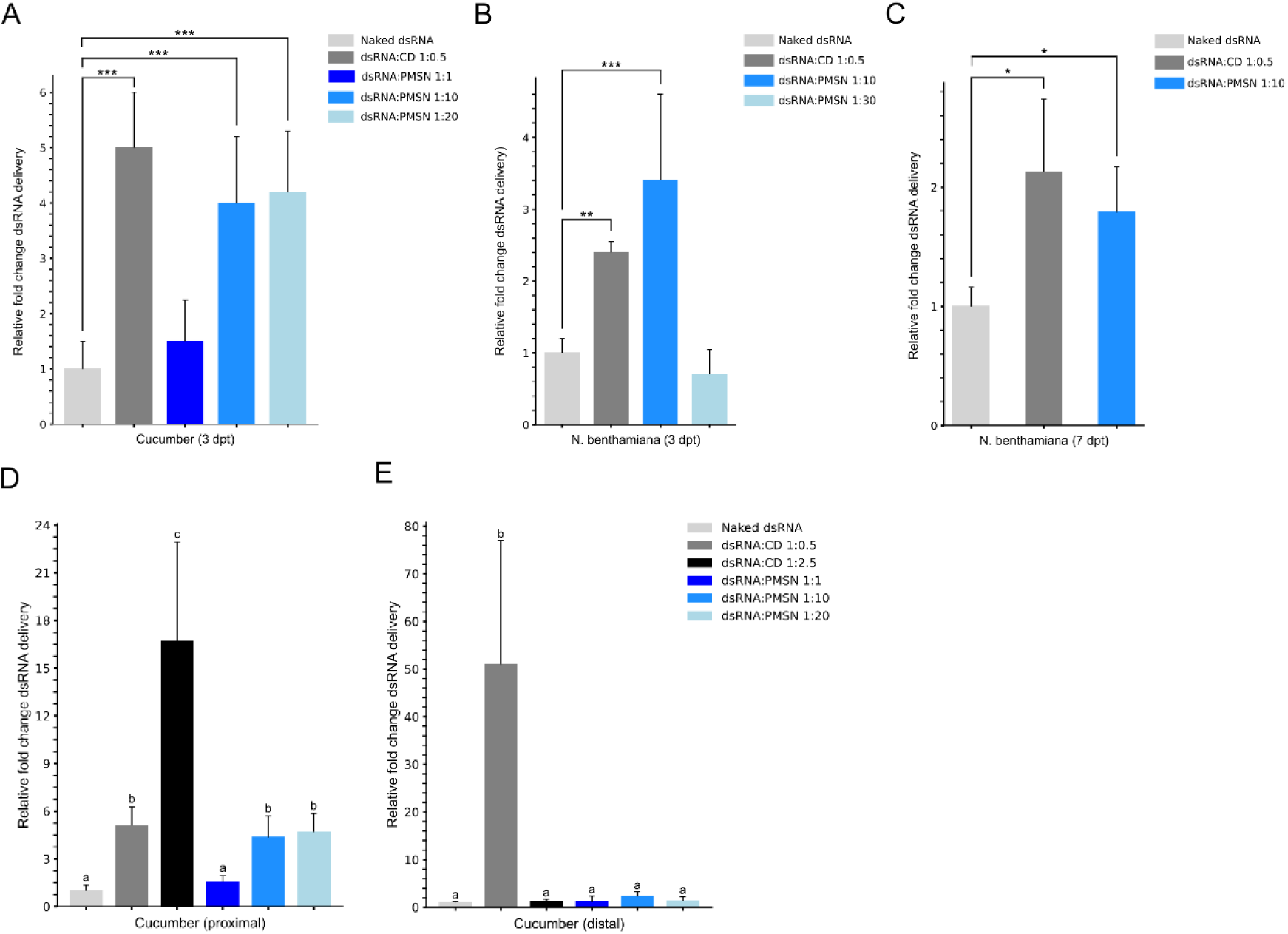
Delivery of dsRNAs to cucumber and *N. benthamiana* plants. A) Relative quantitation in cucumber leaves of dsNbCHE as naked dsRNA or as nanocomposites at different loading ratios after 3 days post treatment (dpt). B) Delivery of dsBCTV in *N. benthamiana* plants after spraying (3 dpt) with naked dsRNA or as dsRNA:NP composites. C) Relative quantitation of dsBCTV in *N. benthamiana* plants after spraying the naked dsRNA or the dsRNA:NP composites at 7 dpt. \**P* < 0.05, \*\**P* < 0.01 and \*\*\**P* < 0.005 represent significance after one-way ANOVA test. Data represent mean ± SEM. D) Relative quantitation in cucumber leaves of dsNbCHE as naked dsRNA or dsRNA:NP composites at different loading ratios after 3 days post treatment (dpt) in the proximal and, E) distal half-leaves. Different letters on the bars refer to significant differences (*P* < 0.05).

**Figure 5.**
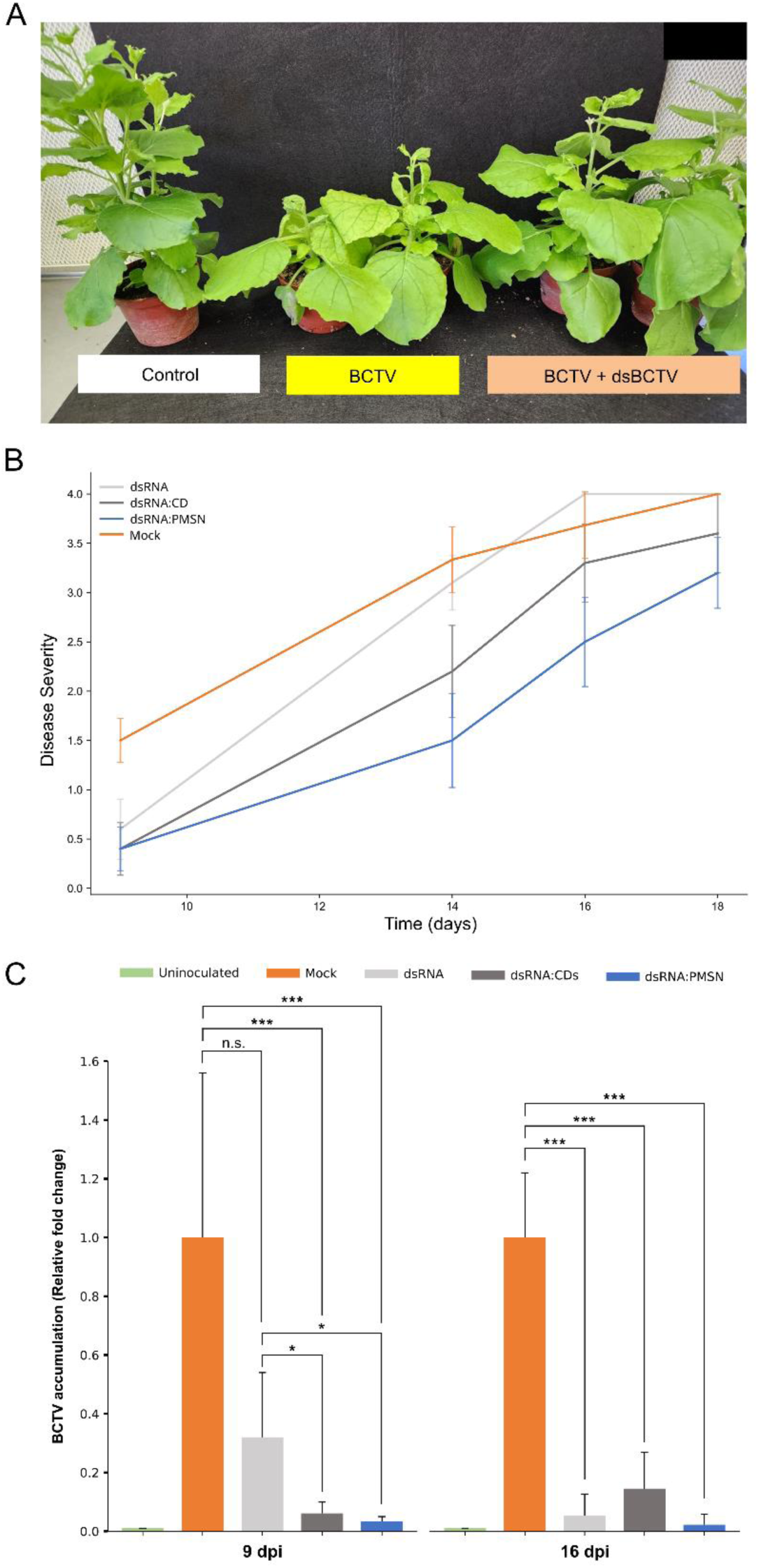
A) Disease symptoms in BCTV-inoculated *N. benthamiana* plants treated with naked dsRNA and the nanocomposites at 14 dpi. B) Disease progress curves for the BCTV-inoculated plants in the different conditions studied. Mock, treated with dsCGMP; dsRNA, treated with naked dsBCTV; dsRNA:CD, treated with dsBCTV coated with CDs; dsRNA:MSNP, treated with the dsBCTV coated with PMSNs. C) Quantitative differences in BCTV accumulation among the different treatments determined at 9- and 16-days post inoculation (dpi). Relative fold differences in BCTV quantitation with respect to the mock control are shown. \**P* < 0.05, \*\**P* < 0.01 and \*\*\**P* < 0.005 represent significance after one-way ANOVA test. Bars represent mean ± SEM.

### 3.4. Evaluation of SIGS treatment against BCTV

In a preliminary experiment, the effect of naked dsRNA targeting BCTV was assessed in plants that were agroinoculated with the virus. Three days after application of naked dsRNA, two leaves from each plant were inoculated with BCTV and symptom development was monitored. In the control, inoculated plants that did not receive dsRNA treatment,, yellowing of the leaves began seven days post-inoculation (dpi). However, in the plants treated with naked dsRNA, symptom onset was delayed until 10 dpi (Fig. 5A). Despite this delay, by 14 dpi, all plants exhibited symptoms with comparable severity, indicating that while naked dsRNA treatment postponed disease onset, it did not ultimately prevent disease progression.

After the preliminary results, a more comprehensive experiment was conducted, including various treatments: naked dsBCTV, nanoparticle-coated dsRNAs (dsBCTV:CD and dsBCTV:PMSN), as well as negative (uninoculated) and positive (mock-treated) controls. Similar to the preliminary trial, BCTV agroinoculation was performed three days after the dsRNA application. Disease progression and viral load were evaluated using symptom scoring and qPCR, respectively. By the 6 dpi, plants in the positive control group began to show initial symptoms of BCTV infection. Symptom severity was recorded at five time points (9, 14, 16, 18, and 23 dpi), following the symptom severity scale described in Table 4. The area under the disease progress curve (AUDPC) was calculated, revealing a statistically significant reduction in disease severity for plants treated with dsBCTV:PMSN compared to untreated inoculated plants (*P* = 0.019) (Table S4, Figure 5B). Regarding virus quantitations, at 9 dpi no significant differences were observed between the mock and the dsRNA treatments at 9 dpi. However, highly significant differences in virus titers resulted when the dsRNAs were delivered with the NPs compared to the mock treatment (Figure 5C). By 16 dpi, the differences compared to the mock were highly significant in all the treatments that showed reduced titers ranging from 8- to 28-fold (Figure 5C).

### 3.5. Evaluation of SIGS treatment against TuMV

The efficacy of dsRNA and nanoparticle-based dsRNA treatments was explored in spray-induced gene silencing (SIGS) context against TuMV. The two TuMV isolates initially considered in this study belong to different clades and pathotypes. Phylogenetic analyses of the nucleotide sequences of the *P1* and *CP* genes allocated the UK1 isolate and DSMZ to the world-B and basal-BR groups, respectively (Figure S3). In preliminary assays, a mix of dsCP and dsHCPro was sprayed on *N. benthamiana* plants followed one hour later by inoculation with the TuMV UK1 isolate. After two weeks, plants that were inoculated with the virus but not dsRNA-treated began to show chlorosis and mosaic symptoms, meanwhile, the sprayed plants did not show clear symptoms of mosaic disease until the end of the assay at 45 dpi (data not shown). In parallel, we observed that symptoms after the inoculation with the DSMZ isolate were more severe. Given that we needed to quantify the disease severity for comparisons among the treatments, we chose the DSMZ isolate for further experiments. Mock-treated TuMV-infected plants displayed strong chlorosis, internode elongation and leaf surface reduction, which was less evident in the dsCP-HCPro or dsCP-HCPro:NP treated plants (Figure 6A). Since chlorosis was a quantifiable parameter, disease severity was assessed through chlorophyll quantification using the SPAD meter. The results indicated that plants treated with either dsCP-HCPro:CD or dsCP-HCPro:PMSN maintained chlorophyll levels comparable to uninoculated controls at both 45 and 65 dpi, indicating effective mitigation of TuMV infection (Figure 6B). In contrast, plants treated with naked dsRNA displayed significantly lower chlorophyll content compared to those receiving nanoparticle-based treatments at 65 dpi. Analysis of viral load across treatments at 7 and 37 dpi showed significant differences between the treated and untreated plants (Figure 6C). At 37 dpi, both dsCP-HCPro:CD and dsCP-HCPro:PMSN treatments significantly reduced viral load compared to the untreated controls, supporting the sustained antiviral effect of nanoparticle-mediated dsRNA delivery over an extended period. At 7 dpi, TuMV-inoculated untreated plants showed an average Cq of 18.7, compared with 20.9, 24.81 and 24.79 of naked dsRNA, dsRNA: PMSN and dsCP-HCPro:CD, respectively. Accordingly, the dsRNA treatments were reduced by 11-fold when compared to no treatments, and the dsCP-HCPro:PMSN and dsCP-HCPro:CD treatments were reduced by 57.8- and 25-fold, respectively. When virus titers were quantified at 37 dpi, the average Cq was 17.7 for the virus-inoculated untreated plants, and 20.1, 21.34 and 22.4 for the naked dsRNA, dsCP-HCPro:PMSN and dsCP-HCPro:CD, respectively. As a result, the viral titer was reduced by 2.4-times in dsRNA-treated plants and 13.5- and 17.3-times, in the dsCP-HCPro:PMSN and dsCP-HCPro:CD-treated plants, respectively.

**Figure 6.**
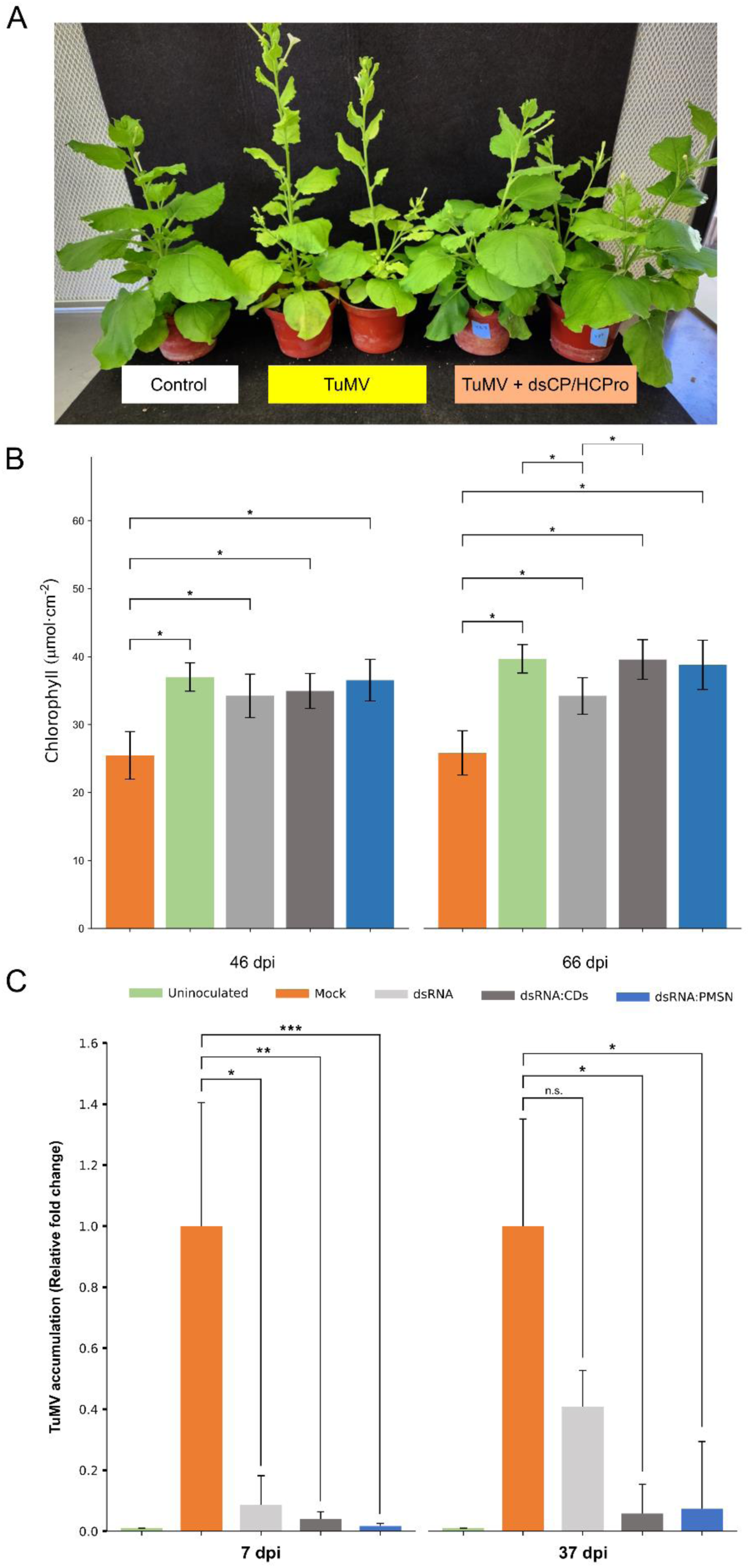
A) Disease symptoms in TuMV-DSMZ-inoculated *N. benthamiana* plants treated with naked dsRNA and the nanocomposites at 25 dpi. The treatments were as follows: Mock, treated with DEPC-treated water; dsRNA, treated with naked dsCP-HCPro; dsRNA:CD, treated with dsCP-HCPro coated with CDs; dsRNA:MSNP, treated with the dsCP-HCPro coated with PMSNs. B) Chlorophyll content in TuMV-inoculated plants in different treatments compared with non-inoculated plants. Determinations were made at 46 and 66 dpi. C) Relative quantitative differences in TuMV accumulation among the treatments with naked and coated dsCP/HCPro determined at 7- and 37-days post inoculation (dpi). \**P* < 0.05, \*\**P* < 0.01 and \*\*\**P* < 0.005 represent significance after one-way ANOVA test. The effect of the treatment with naked dsCP/HC-Pro at 37 dpi resulted non-significant (*P* = 0.062). Bars represent mean ± SEM.

## 4. Discussion

In this study, we have investigated SIGS-derived resistance to BCTV and TuMV in *N. benthamiana*. To improve dsRNA delivery to plants, two nanoparticle formulations CDs and PMSNs were used. The CDs used in this study were synthesized following Delgado-Martín et al. (2022a) and displayed similar properties such as small size (< 5 nm), high photoluminescence, and capacity to bind to dsRNA. The successful synthesis and functionalization of PMSNs were confirmed through a combination of TEM, DLS, and FT-IR, ensuring that the nanoparticles were suitable for dsRNA binding and delivery. The average diameter of PMSNs as determined by DLS differed from the size characterization of the PMSNs by TEM. In DLS, the hydrodynamic diameter of nanoparticles dispersed in liquids is generally reported to be larger than their actual diameter due to the presence of other molecules such as water molecules around the particles. In fact, the more hydrophilic the particles are, the more water molecules are attracted to them and their hydrodynamic diameter increases (Behzadi et al. 2018). The loading of nucleic acid molecules on nanoparticles with positive charges occurs through several forces, including the electrostatic attraction between the negative charge in the structure of the nucleic acids and the positive charge on the nanoparticles, free electrolytes in the solution, and hydrogen bonding between the ssRNA and the surface of the nanoparticles (Della Rosa et al. 2020). Functionalizing the surface of silica nanoparticles with cationic agents including aminopropyl groups such as aminopropyltrimethoxysilane (Gao et al. 2009; Cai et al. 2024), large polycations such as PEI (Ufer et al. 2008), poly-L-lysine (PLL) (Yu-Qin et al. 2012), diethanolamine (DEA), polyamine compounds such as spermidine (Cho et al. 2023) or triethanolamine (TEA) (Sangwan et al. 2023) by covering negative silanol groups with positive amine groups, leads to increasing electrostatic interactions between the surfaces of MSNs and nucleic acids (Zolghadrnasab et al. 2021). In our study, the electrostatic attraction between the phosphate groups present in the ribonucleic acid structure of the dsRNA molecules and the positive charge of the amine polymer groups present on the PMSNs was used in the loading stage. Although FT-IR spectroscopy can be used to detect sugars and phosphates of nucleic acids from their characteristic absorption bands (Banyay et al. 2003; Nandiyanto et al. 2019), in our nanocomposite we could clearly detect only one of them, probably because of the low proportion of dsRNA with respect to PMSNs in the preparations that resulted in the masking of most of the specific peaks. The increase in the hydrodynamic diameter and the negative value of the ζ potential (charge) of the dsRNA:PMSNs complex showed that the dsRNA molecules covered the surface of the polyamine mesoporous silica nanoparticles with a positive charge and reduced the overall charge of the complex to −10 mV. Similar results of siRNA loading on MSNs have been reported after the MSNs were functionalized with amine, that increased from −10.8 to +16.2 and decreased to −15.2 mV after loading with siRNA (Heidari et al. 2021).

We observed that increasing relative the amount of PMSNs compared to dsRNA increased the loading efficiency up to 69% in a ratio of 1:40. By lowering the PMSNs ratio to 1:10, a significant increase in their loading capacity was observed compared to the 1:20 ratio, suggesting a gradual saturation of the nanoparticle surface with dsRNA. In parallel, when decreasing the PMSN ratios from 1:10 to 1:5 and 1:1, a noticeable decrease in the upward trend (the slope of the graph) of the loading capacity was observed; and inversely, the loading efficiency decreased from 42% to 22.8% and 4.5%, respectively, as also evidenced in retardation agarose gels. Similarly, the maximum loading efficiency of dsRNA_AC2_ from the ToLCNDV genome on MSNs of different sizes amino-MSNP_DEA_, amino-MSNP_TEA_ and amino-MSNP_NH3_ was reported as 46%, 65% and 71%, respectively. Also, in the same report, it has been shown that amino-MSNP_DEA_ (∼10 nm, 4 ± 0.2 mV), amino-MSNP_TEA_ (∼32 nm, 4.5 ± 0.2 mV), amino-MSNP_NH3_ (∼66 nm, 5 ± 0.1 mV) whose positive charges were derived from APTES, can enter the *N. benthamiana* leaf and root cells by being rubbed and soaked, respectively (Sangwan et al. 2023). In our work, the maximum loading capacity was calculated in the 1:1 ratio of dsRNA:PMSNs as 45.9 ng·μg⁻¹. These results were consistent with the results of another study that used PMSNs to transfer plasmid DNA (pDNA) containing the GUS gene expression construct to callus cells of tobacco plants, and the maximum loading capacity reported in a 1:1 ratio of pDNA loading on PMSNs was 43.4 ng·μg⁻¹ (Zolghadrnasab et al. 2021). Recently, it was shown that the capacity of loading siRNAs was conversely proportional to MSN diameter (Cai et al. 2024).

The fitting of the dsRNA:PMSNs adsorption isotherm on the common models predicting the surface adsorption behavior of nanoparticles showed that the Langmuir-Freundlich model can provide a more accurate description of the interaction between dsRNA and PMSNs than the other two models. In the Langmuir model, the uniform distribution of active groups on the surface of nanoparticles is assumed and there is no lateral interaction between particles (Miedaner et al. 2006). This model is valid and usable when monolayer absorption occurs at the level of nanoparticles; absorption includes a certain number of identical areas with a single loading method and it is not possible to move the adsorbed substance on the absorbent surface (Gautam et al. 2014). The Freundlich model is a type of adsorption model in which the adsorbed substance forms a layer on the absorbent surface (Singh 2016), but unlike the Langmuir model, multilayer adsorption and interaction between the adsorbed substances are possible in this model (Al-Ghouti and Da’ana 2020). The Freundlich model refers to the non-uniformity of the surface of the molecules and the exponential distribution of the active regions and their energy (Gast et al. 1997). By integrating two Langmuir and Freundlich models, it is possible to simultaneously simulate Langmuir and Freundlich absorbing and absorbing behavior (Jeppu and Clement 2012). In low and high adsorbed concentrations, the Langmuir-Freundlich adsorption isotherm model shows the same behavior as the Freundlich adsorption isotherm model and the Langmuir adsorption isotherm model, respectively (Ayawei et al. 2017).

High concentrations of NaCl and EDTA lead to the release of RNA from a column made of mesoporous silica particles attached to spermidine polyamine (Cho et al. 2023). However, treatment of the dsRNA:PMSNs composite with EDTA did not result in dsRNA release. DsRNA molecules present more phosphate groups than ssRNA due to their duality, and as a result, stronger electrostatic interactions between dsRNA and PMSNs are created. Hence, it seems that higher concentrations of chelating agents are required to disrupt stronger bindings between dsRNA and PMSNs and maximal release of dsRNA (Cho et al. 2023). SDS is a well-known anionic surfactant that on behalf of its sulfate group is strongly attached to the positively charged areas of PMSNs (Hansen et al. 2009). Following the reduction of the positive charge or the neutralization of the surface charge of nanoparticles mediated by SDS, the electrostatic interaction between dsRNA and PMSNs was lost, allowing the release of the molecules.

The NP- assisted delivery on plants aims to enhance the delivery and stability of dsRNAs in plant tissues. Here, we used CDs and PMSNs both functionalized with positive amine groups derived from PEI on their surfaces to increase the availability of the plant cells to dsRNA molecules and eventually improve the SIGS efficiency. Among the different application methods of exogenous dsRNA, spray seems the most accessible, especially in large-scale usage which allows the farmers to apply the dsRNA in the form of particles (Mat Jalaluddin et al. 2023; Koeppe et al. 2023). CDs and MSNs have been shown to increase the silencing of endogenous plant genes when delivering ds- or siRNAs (Schwartz et al. 2020; Cai et al. 2024). The delivery of dsRNA was reported to be enhanced when mediated by CDs in *N. benthamiana* and cucumber and resulted in the increase of derived siRNAs (Delgado-Martín et al. 2022a). The efficiency of dsRNA molecules can be increased by NPs’ ability to shield them from intracellular and environmental degradation, extending their half-life (Parker et al. 2019; Bachman et al. 2020). For example, when dsRNA and siRNA were loaded onto LDH nanoparticles (Mitter et al. 2017a) or MSNs (Parnian et al. 2022), respectively, the dsRNA molecules were not affected by RNase A treatments. However, CDs were unable to protect dsRNA from RNase A nuclease digestion under unsaturated conditions (Delgado-Martín et al. 2022a). In our conditions, we have not observed protection from RNase A of dsRNA adsorbed to PMSNs (not shown), but have focused, instead, on the capability of nanoparticles in improving the release and stability of dsRNAs in plant leaves. Quantitative RT-PCR analysis at three days post-application revealed that both CDs and PMSNs significantly improved dsRNA delivery into cucumber and *N. benthamiana* leaves compared to naked dsRNAs. In cucumber plants treated with a 1:10 ratio of dsRNA:PMSN composites, there was a significant increase in dsRNA accumulation compared to naked treatments. Similarly, in *N. benthamiana*, a 3.4-fold increase in dsRNA accumulation was observed with a 1:10 ratio compared to naked treatments. These results are consistent with previous studies showing that nanoparticle-mediated delivery systems can enhance the uptake and stability of exogenous nucleic acids in plants (Jiang et al. 2014; Schwartz et al. 2020; Delgado-Martín et al. 2022a). The stability of delivered dsRNAs over time was also assessed by qRT-PCR at seven days post-application. Both CDs and PMSNs significantly enhanced the persistence of dsRNAs within plant tissues compared to naked-dsRNA treatments. DsRNA movement in the plant is one of the current limitations of SIGS. In our experiments, we observed that dsRNA movement is restricted except when the dsRNA is delivered in the form of dsRNA:CD nanocomposite in the 1:0.5 ratio, confirming the results reported elsewhere (Delgado-Martín et al. 2022b).

Since viral replication cycles often extend beyond initial treatment periods, the increase in delivery and stability mediated by NPs is desirable (Mitter et al. 2017a; Dalakouras et al. 2020; Xu et al. 2023; Sangwan et al. 2023). Most of the studies reported to date on the use of exogenous dsRNA against plant viruses have been performed on RNA viruses, given that up to our knowledge, effective control for DNA viruses with this method is scarce, as has been reported for the begomoviruses tomato leaf curl virus (ToLCV), tomato yellow leaf curl virus (TYLCV) and ToLCNDV (Namgial et al. 2019; Melita et al. 2021; Sangwan et al. 2023; Frascati et al. 2024). In general, the infection method for the virus was agroinoculation into the abaxial side of the leaf, followed by the simultaneous application of dsRNA to the adaxial surface of the same leaf. However, it has been recently shown that dsRNA delivered by amine-functionalized MSNPs could decrease viral titers in ToLCNDV-infected *N. benthamiana* plants when supplied to roots or leaves (Sangwan et al. 2023). In contrast, when another DNA virus, as is the case for the begomovirus tomato severe rugose virus (ToSRV), was inoculated by its insect vector, the whitefly, that directly transfers the virus to the vascular tissue of the plant, the dsRNA treatment was unsuccessful (Rego-Machado et al. 2020). In this case, the failure of the SIGS method in generating resistance may be due to the possibility that the virus enters the vascular tissue before being inhibited by the interfering RNA induced by dsRNAs in plant leaves (Rêgo-Machado et al. 2023). In a recent work, it has been shown that dsRNA derived from the *AC2* gene, when coated with MSNPs, produced amelioration of symptoms and decreased virus titers after infection with ToLCNDV (Sangwan et al. 2023). The same BCTV gene segment chosen in our study for SIGS were previously used by Montazeri and col. (2024) to transiently express the viral sense, anti-sense, and hairpin in *N. benthamiana* and sugar beet to confer the resistance against BCTV and BCTIV. They observed that the hairpin form of RNA could significantly delay the disease symptom appearance compared to non-transgenic *N. benthamiana*, as well as the transgenic *N. benthamiana* transformed with sense and anti-sense constructs. In our preliminary assays, when using naked dsRNAs targeting agroinoculated BCTV, we observed that symptom onset was delayed by three days compared to untreated controls. However, by 14-dpi, all plants exhibited symptoms with comparable severity, indicating that while naked dsRNA treatment could delay disease progression slightly through SIGS, it was insufficient for long-term protection against BCTV infection. Noteworthy, the use of nanoparticle-coated dsRNAs showed more interesting results. In particular, plants treated with dsRNA:PMSN formulations exhibited a statistically significant reduction in disease severity compared to untreated controls and those treated with naked dsRNA. Although at 7-dpi, no significant differences in viral load were observed between treatments, by 14-dpi, plants treated with dsRNA:PMSN exhibited a two-fold reduction in viral load compared to untreated controls. Even though the SIGS method did not provide the plants with a desired long resistance against this DNA virus, it is conceivable that multiple applications of the NP-dsRNAs could boost their effectiveness (Cai et al. 2024).

According to the literature, the induction of resistance by the exogenous application of dsRNA seems to be more efficient for RNA viruses (Tenllado and Díaz-Ruíz 2001; Yin et al. 2009; Delgado-Martín et al. 2022b). The 3′ end of the genome of potyviruses such as TuMV is a highly conserved region that includes the *NIa*, *NIb* and *CP* genes (Yang et al. 2021). Several reports have shown that with external application of exogenous dsRNA from *CP* and *NIb* genes, dsRNA_CP_ had better efficiency in reducing potyvirus infection (Sun et al. 2010; Worrall et al. 2019). The HC-Pro protein encoded in the TuMV genome is an example of a multifunctional protein that carries at least three different functions: plant-to-plant transmission of the virus, polyprotein maturation, and suppression of the RNA silencing process (Valli et al. 2018). Local application of dsRNA_HC-Pro_ with different lengths from 500 to 1100 bp has significantly reduced the infection caused by the potyviruses TEV (*Potyvirus nicotianainsculpentis*), PPV (*Potyvirus plumpoxi*), PVY (*Potyvirus yituberosi*), ZYMV (*Potyvirus cucurbitaflavitesselati*), PRSV (*Potyvirus papayanuli*) or PepMoV (*Potyvirus capsimaculae*) (Tenllado and Díaz-Ruíz 2001; Sun et al. 2010; Kaldis et al. 2018; Yoon et al. 2021). Consequently, we used combined CP+HC-Pro TuMV-derived dsRNAs in our research. Recently, a reduction of symptoms in tomato, pepper and tobacco by 60% and a decrease of virus titers mediated by nanoparticle (including CDs)-coated dsRNAs have been reported in potato virus Y-infected plants (Xu et al. 2023). In the *N. benthamiana* plants treated with either dsRNA:CD or dsRNA:PMSN we observed chlorophyll levels comparable to uninoculated controls at both 45 and 65 dpi, indicating effective mitigation of TuMV infection. In contrast, plants treated with naked dsRNA displayed significantly lower chlorophyll content compared to those receiving nanoparticle-based treatments at 65 dpi. Analysis of viral load across treatments at 7 and 33 dpi showed significant differences between treated and untreated plants. At 7 dpi, TuMV-inoculated untreated plants showed lower Cq values than plants treated with naked dsRNA. Furthermore, plants treated with dsRNA:PMSN or dsRNA:CD exhibited even higher Cq values than the inoculated plants treated with naked dsRNA only. This corresponds to an approximate reduction in viral load by 57.8- and 25- fold for dsRNA:PMSN and dsRNA:CD treatments compared to untreated controls. At 33 dpi, both nanoparticle-based treatments continued to show significant reductions in viral load compared to untreated controls, although the differences with the treatments with naked dsRNA were negligible. It is then possible that after long periods the increase of dsRNA delivery mediated by nanoparticles is no longer effective, at least for this potyvirus.

In this work, we have shown that the use of carbon dots and mesoporous silica nanoparticles as carriers for delivering exogenous dsRNAs represents a plausible strategy for controlling DNA and RNA plant viruses through RNAi. Nonetheless, several challenges remain before this kind of nanoparticle-mediated RNAi technology can be widely adopted for crop protection. Indeed, the environmental fate and dissipation kinetics of nanoparticle-coated dsRNAs need further investigation to ensure their safety for non-target organisms and ecosystems (Bachman et al. 2020). While some research has demonstrated that unformulated dsRNAs break down quickly in soil (Dubelman et al. 2014), formulations of nanoparticles intended to improve uptake or stability may alter their degradation kinetics. A further obstacle is the expense of manufacturing large amounts of dsRNAs for use in commercial settings, in particular for agricultural applications. A significant challenge is the high cost of producing large quantities of dsRNAs for use in commercial settings, particularly in agricultural applications (Dalakouras et al. 2020). Microbial-based production systems for generating cost-effective sources of dsRNAs may help address this issue (Ahn et al. 2019; Ma et al. 2020; Delgado-Martín and Velasco 2021). Finally, regulatory frameworks for assessing the environmental risks associated with nanoparticle-mediated RNAi products are still evolving while comprehensive risk assessments will be necessary to ensure the safe use of dsRNA and nanoparticles in agriculture.

## Acknowledgements

This work was funded by grant PID2021-125787OR-C32 from the AEI/MICIN of Spain. This work has been supported by the Center for International Scientific Studies & Collaboration (CISSC), Ministry of Science, Research and Technology of Iran.

## Notes

### Competing Interest Statement

The authors have declared no competing interest.

