## Supporting information for "Topical application of carbon dots and mesoporous silica nanoparticle-derived dsRNA-nanocomposites for the control of beet curly top virus and turnip mosaic virus"

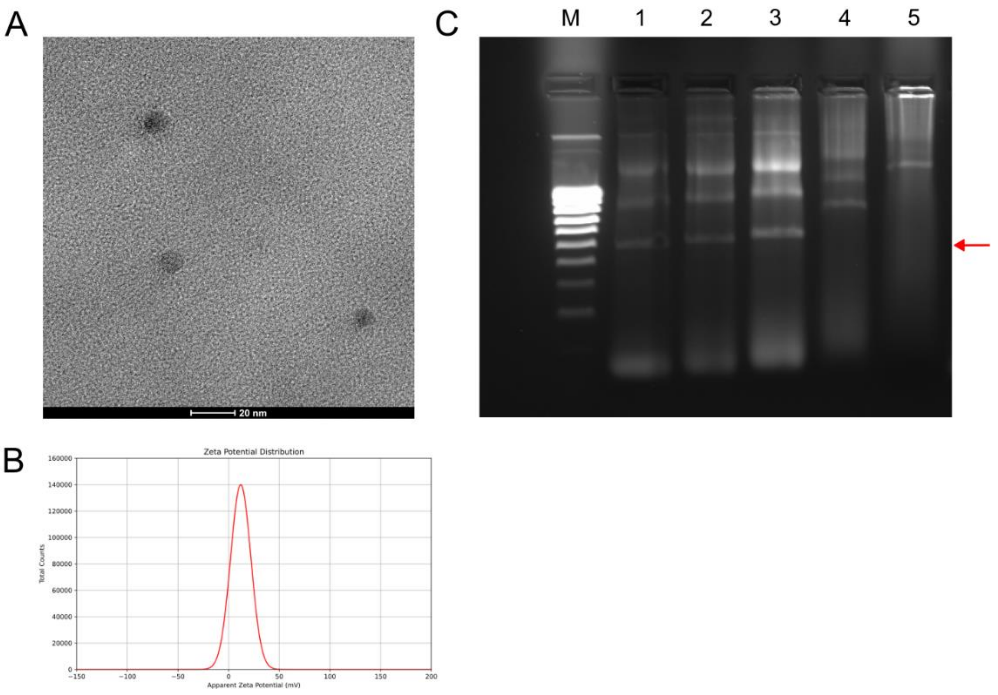


Figure S1. TEM image (A) and ζ potential (C) as measured with the Zetasizer of the carbon dots obtained in this work. B) Retardation gel of dsRNA and dsRNA:CD composites. M. NZYDNA Ladder V, (1) naked dsRNA, (2) dsRNA:CD 1:0.1, (3) dsRNA:CD 1:0.2, (4) dsRNA:CD 1:0.5, (5) dsRNA:CD 1:1. The arrow points to the dsRNA band of the RNA bacterial extraction in the first lane. Nucleic acids bands result retarded in the electrophoresis with the increasing amounts of carbon dots.

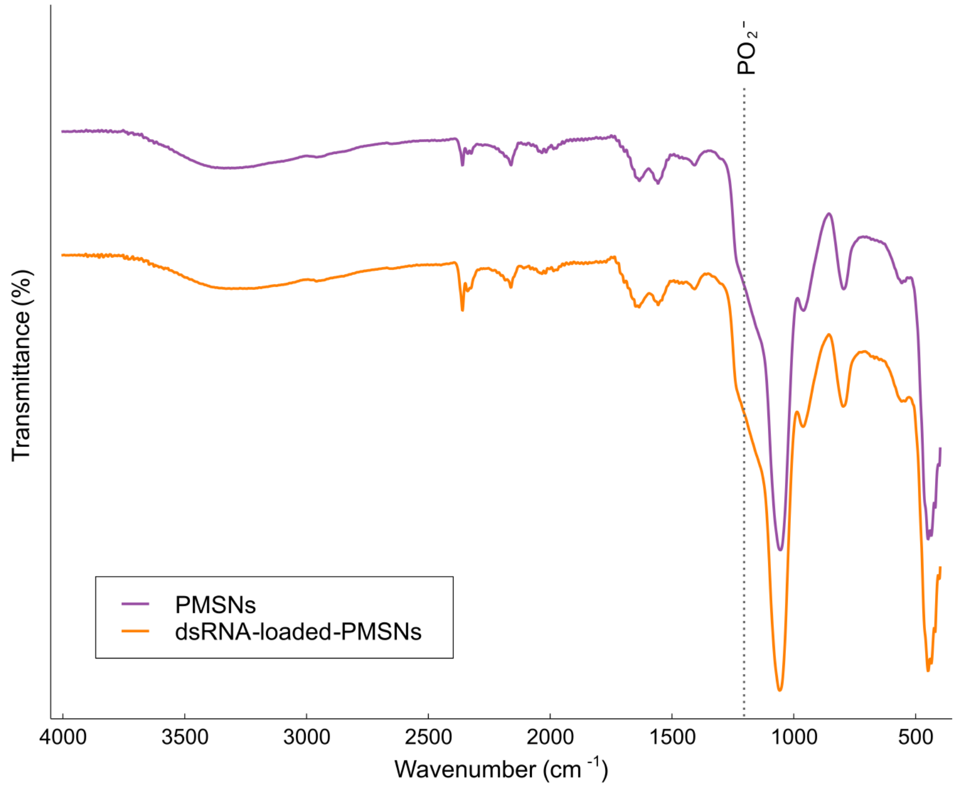


Figure S2. FT-IR spectra of the polyamine mesoporous silica nanoparticles (PMSNs) and the dsRNA:PMSNs composites.

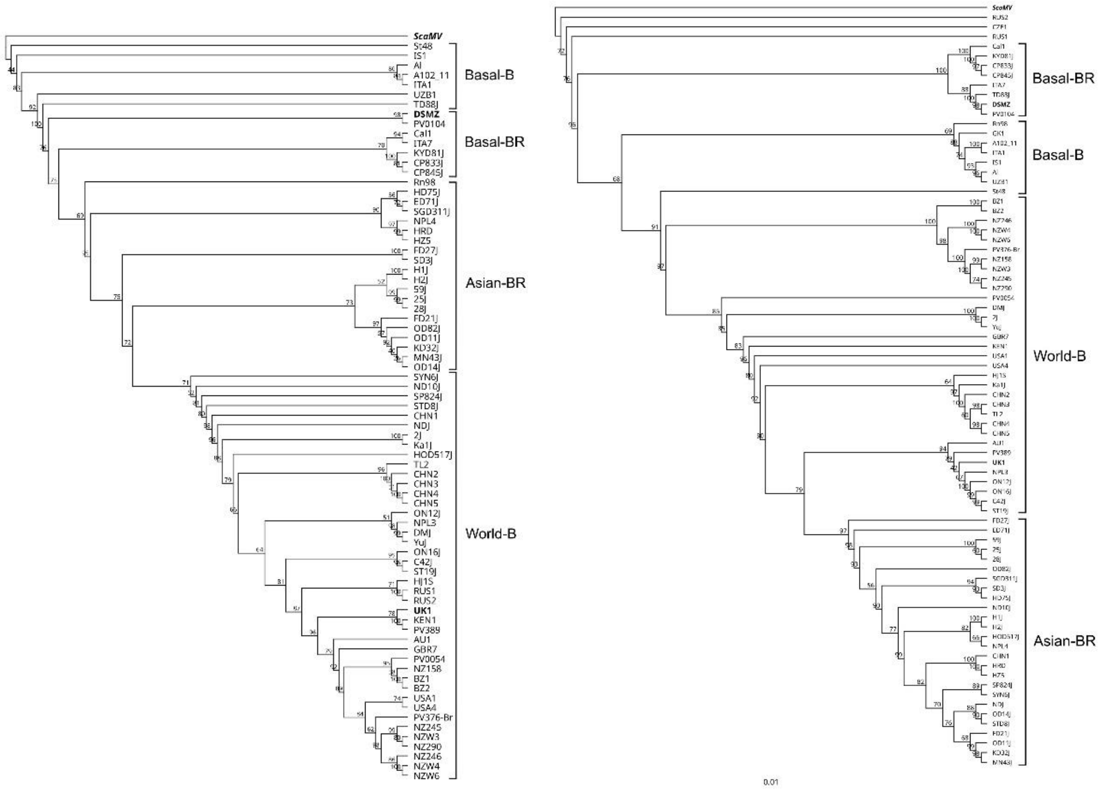


Figure S3. Phylogenetic trees obtained for the CP (left) and P1 (right) regions of 74 isolates of TuMV described elsewhere (Ohshima et al., 2002), in addition to the DSMZ and UK1 isolates (in bold). Scallion mosaic virus (ScMV), a member of the TuMV phylogenetic group, was used as outgroup taxon. The trees were generated after MAFFT alignment in the IQ-TREE software using the default parameters for nucleotide sequence analyses. The best-fit models according to BIC were TIM+F+G4 and TIM2+F+G4 for the CP and P1 segments, respectively.

Table S1. Primers used in this work.

| Name | Target/purpose | Sequence 5'-3' | Reference |
|---|---|---|---|
| TuMV_UKCP_L4440_F | CP gene of TuMV UK1 (In-Fusion cloning to L4440) | CGGTATCGATAAGCTGCAGAAGCAGTTGGCACTCA | This work |
| TuMV_UKCP_L4440_R | " | GGAGACCGGCAGATCTCTCTCGCACGTATTGGAGT | " |
| TuMV_UKHC1_L4440_F | HCPro gene of TuMV UK1 (In-Fusion cloning to L4440) | CGGTATCGATAAGCTTCACAAGGTGATTTTGTCCATGC | " |
| TuMV_UKHC_L4440_R | " | GGAGACCGGCAGATCTTTCAAGCTCGACTCCAAGTTGC | " |
| TuMV_DSCP_L4440_F | CP gene of TuMV DSMZ (In-Fusion cloning to L4440) | CGGTATCGATAAGCTGCAGAAGCAGTTGGCACTCA | " |
| TuMV_DSCP_L4440_R | " | GGAGACCGGCAGATCTCTCTCGCACGTATTGGAGT | " |
| TuMV_DSHC_L4440F | HCPro gene of TuMV DSMZ (In-Fusion cloning to L4440) | CGGTATCGATAAGCTTTGCAAGGCGACTTTGTTCATGC | " |
| TuMV_DSHC_L4440R | " | GGAGACCGGCAGATCTTTCAAACTCGATTCCAAGTTAC | " |
| qTuMVDSH215F | HCPro gene of TuMV DSMZ (qPCR) | ATGTCAAGGAGTCGCAAGCA | “ |
| qTuMVDSH354R | " | GTTGGCAACATCCGGGTAGA | " |
| qTuMUKH586F | HCPro gene of TuMV UK1 (qPCR) | ATGTGTGACAACCAGCTCGAT | " |
| qTuMUKH750R | " | GATTGCAAGTTTCCGTGACCC | " |
| qBCTV-F | Beet severe curly top virus V1 gene (qPCR) | CAGGCTATGCCGTCTACTTATC | " |
| qBCTV-R | " | TTCCCTCCATATCCAGTACCA | " |
| attB1NbCH1883 | *N. benthamiana* Magnesium chelatase subunit ChlH (XM_004149349) (Gateway cloning) | GGGGACAAGTTTGTACAAAAAAGCAGGCTATGAGGGTGACCCGATGAGA | " |
| attB2NbCH2473 | " | GGGGACCACTTTGTACAAGAAAGCTGGGTCTTCCATCGCTGTTGGAGGT | " |
| qNbCHE-F | *Nb* Magnesium chelatase quantitation (qPCR) | TTCCTACCACAAGGTCACGC | " |
| qNbCHE-R | " | TTGAAAGACTCAGGCCGTGG | " |
| Cs18S-F | *Cucumis sativus* RNA 18S gene (Cs reference gene in RT-qPCR) | GGCGGATGTTGCTTTAAGGA | Gil-Salas et al., 2009 |
| Cs18S-R | | GTGGTGCCCTTCCGTCAAT | " |
| NbEF-1α-F | Nb Elongation factor 1α (EF1a) (qPCR reference gene) | AGCTTTACCTCCCAAGTCATC | Liu et al., 2012 |
| NbEF-1α-R | " | AGAACGCCTGTCAATCTTGG | " |

Table S2. Rating scale for the beet curly top disease symptoms used in this study.

| Score | Description of symptoms |
|---|---|
| 0 | Healthy, no symptoms |
| 1 | Vein clearing of leaves, slight enation of veins on the abaxial part of the leaves, leaves yellowing |
| 2 | Slight leaf curling of the edges of the new leaves, enation of veins on the abaxial part of the leaves |
| 3 | Leaf curling of the edges of the new leaves, no or slight stunting |
| 4 | Severe curling of the central leaves, moderate or severe stunting, plant dwarfism |

Table S3. Langmuir, Freundlich and Langmuir-Freundlich fitting parameters for the adsorption of dsRNA into the PMSNs at 1:10 ratio.

| Model | Constants | | $R^2$ |
|---|---|---|---|
| Langmuir | $q_{max}$ | 151.84 | 0.893 |
| | $K_L$ | 0.0041 | |
| Freundlich | $K_F$ | 1.17 | 0.877 |
| | n | 1.27 | |
| Langmuir-Freundlich | $q_{max}$ | 51.66 | 0.950 |
| | $K_{LF}$ | 0.0200 | |
| | n | 2.75 | |

Table S4. Statistical analysis of AUDPCs from the BCTV-inoculated plants.

| Condition | Mean AUDPC | SEM |
|---|---|---|
| dsBCTV | 24.35 | 1.52 |
| dsBCTV:CD | 18.90 | 2.77 |
| dsBCTV:PMSN | 14.45 | 2.73 |
| Mock | 26.78 | 2.01 |
| Shapiro-Wilk test p-value | 0.0000 | |
| Levene's test p-value | 0.1462 | |
| Test used | Kruskal-Wallis | |
| H-statistic | 13.09 | |
| *P*-value | 0.0045 | |

| Group 1 | Group 2 | *P*-value |
|---|---|---|
| Mock | dsBCTV | 0.9269 |
| Mock | dsBCTV:CD | 0.2134 |
| Mock | dsBCTV:PMSN | 0.0194 |
| dsBCTV | dsBCTV:CD | 0.3998 |
| dsBCTV | dsBCTV:PMSN | 0.0335 |
| dsBCTV:CD | dsBCTV:PMSN | 0.5720 |

Tukey's HSD Post Hoc Analysis
